# Left atrial flow and thrombosis risk from 4D CT contrast dynamics by physics-informed neural network and indicator dilution theory

**DOI:** 10.64898/2026.03.31.715623

**Authors:** Bahetihazi Maidu, Alejandro Gonzalo, Manuel Guerrero-Hurtado, Clarissa Bargellini, Pablo Martinez-Legazpi, Javier Bermejo, Francisco Contijoch, Oscar Flores, Manuel García-Villalba, Elliot McVeigh, Andrew M. Kahn, Juan C. del Alamo

## Abstract

Atrial fibrillation (AF) promotes blood stasis and thrombus formation, most often within the left atrial appendage (LAA), and can lead to stroke or transient ischemic attack (TIA). Time-resolved contrast-enhanced computed tomography (4D CT) captures left atrial (LA) opacification and washout, but it does not directly provide quantitative stasis metrics such as blood residence time. Patient-specific computational fluid dynamics (CFD) can quantify LA/LAA residence time, yet routine clinical use is limited by computational cost and sensitivity to patient-specific boundary conditions. Here, we present two complementary approaches to infer time-resolved 3D residence time fields directly from contrast dynamics. First, a physics-informed neural network (PINN) treats contrast as a passive scalar and jointly reconstructs velocity and residence time by enforcing the incompressible Navier–Stokes equations and transport equations for contrast concentration and residence time in moving, patient-specific LA anatomies. Second, an indicator dilution theory (IDT) formulation computes voxelwise, time-resolved residence time maps from contrast time curves alone by constructing a PV-referenced impulse response and modeling transport with a tank-in-series model with spatially dependent parameters. Both methods are benchmarked against patient-specific CFD in six cases spanning diverse LA function, including three patients with TIA or thrombus in the LAA and three patients free of events. Both approaches reproduce expected spatial and temporal trends, with higher residence time in the distal LAA and higher LAA residence time in cases with TIA or thrombus. IDT demonstrates the closest agreement with CFD across the full range of residence times and produces maps in seconds, facilitating clinical translation. In contrast, the PINN additionally recovers phase-dependent atrial flow structures, but tends to smooth and underestimate the highest residence-time regions and requires hours of training. Together, these results support a scalable workflow in which IDT enables rapid stasis screening from contrast CT, and PINNs provide a complementary pathway for detailed, patient-specific hemodynamic inference when full-field flow information is needed.

## 1 Introduction

During atrial fibrillation (AF), the left atrium (LA) contracts weakly and irregularly, slowing down blood flow in regions like the left atrial appendage (LAA). These conditions promote the formation of thrombi, some of which may travel to the brain and cause ischemic strokes. Consequently, 15% of all strokes are thought to be caused by AF [1]. These strokes are associated with worse outcomes, more disability, and higher mortality than ischemic strokes in the absence of AF [2]. Anticoagulation therapies for stroke prevention are based on clinical evidence that their benefits statistically outweigh their risks in large groups of AF patients with certain demographic and comorbid factors [3]. These factors are usually aggregated into numerical scores (e.g., CHA_2_DS_2_-VASc) that, when applied to individual patients, have modest accuracy [4]. Currently, there are no quantitative tools to personalize stroke risk prediction in AF patients.

Given that the vast majority of LA thrombi form inside the LAA [5], estimating LAA thrombosis risk is considered comparable to estimating stroke risk during AF. From a mechanistic standpoint, prolonged blood residence time in the LAA is an appealing biomarker because it directly quantifies the slow flow conditions that favor thrombogenesis. Slow or reduced swirling blood flow inside the LA, measured by 4D flow magnetic resonance imaging (MRI) [6–8], has been associated with AF and ischemic stroke. However, current 4D flow MRI methods cannot resolve flow inside the small-size LAA, especially its distal part. Echocardiographic vector flow mapping (VFM) detects slow flow regions to predict the risk of left ventricular (LV) thrombosis and associated brain embolisms [9–11]. Applying VFM to the LAA would require transesophageal echocardiography, which is unlikely to be adopted to risk-stratify AF patients due to its invasive nature.

Computed tomography (CT) produces high-resolution images with quick acquisitions. While no methods are yet available to measure blood flow from CT, it is recognized that slow flow can create noticeable contrast gradients in the LAA. In particular, poor CT enhancement in the distal LAA has been associated with stroke risk in AF patients [12, 13]. However, the temporal manifestation and spatial steepness of these gradients are likely to depend on each individual’s LA and LAA filling dynamics. Consequently, CT studies using different imaging and analysis protocols have not reached consensus about the relationship between LAA contrast gradient and stroke risk [14]. This limitation suggests that qualitative assessment of LAA opacification alone may not provide a robust patient-specific marker of stasis. However, because iodinated contrast behaves approximately as a passively advected scalar, its spatiotemporal dynamics still encode information about intracardiac transport, motivating quantitative approaches that infer residence time from contrast time curves while accounting for the governing flow physics.

Computational fluid dynamics (CFD) analysis is garnering attention since it can finely resolve stagnant regions inside each patient’s LAA from CT image segmentation [15–26]. When parameterized correctly, CFD models can compute essentially exact flow fields by representing the governing equations of flow physics. However, CFD parameterization is sensitive to assumptions about LA wall motion [27], blood rheology [28], and inflow boundary conditions [29]. Furthermore, creating CFD meshes, selecting numerical discretization schemes, and setting CFD resolution to balance accuracy with compute time all require significant technical expertise [30]. Machine learning and, in particular, deep learning (DL) offers a versatile alternative to CFD for finding complex relationships between cardiac chamber anatomy, blood flow, and clotting risk [31–33]. However, these approaches are often constrained by the requirement for extensive, high-quality labeled biomedical imaging training data, which limits their clinical applicability [34].

Physics-informed neural networks (PINNs) combine patient-specific data with governing equations to infer hidden flow information, with a paradigmatic example being flow reconstruction in a cerebral aneurysm from contrast dynamics [35, 36]. Recent developments in low-dose multi-cycle 4D contrast CT enable imaging of LA/LAA filling and emptying over multiple heartbeats [37], creating an opportunity to apply PINNs to intracardiac transport. In this context, PINNs provide a framework to infer contrast concentration, flow and residence time fields from spatiotemporal contrast data. Patient-specific training minimizes a loss function that includes a data-fidelity term for contrast concentration, the residuals of the Navier–Stokes equations, and transport equations for both contrast concentration and residence time. Notably, unlike conventional CFD workflows, such a framework does not require an explicit computational mesh of the LA or complete explicit prescription of flow boundary conditions during inference.

Indicator dilution theory (IDT) has a long clinical history for estimating blood flow parameters, vascular volume, and cardiac output following bolus injection of a tracer [38]. Its simplicity and rapid inference capabilities have motivated extensive applications across medical imaging modalities, including ultrasound [39], MRI [40], and CT [41]. In this setting, IDT can be used to infer residence time from contrast time curves by modeling chamber transport as a linear response system with a parameterized transport function (e.g., a gamma-variate function) [42].

In this study, we use CFD-generated contrast dynamics as ground truth to investigate two complementary approaches for inferring LA/LAA residence time from contrast CT: (i) a PINN that reconstructs velocity, contrast concentration, and residence time fields by enforcing the incompressible Navier–Stokes equations together with transport equations for contrast and residence time, and (ii) an IDT framework that computes voxelwise residence time directly from temporal contrast curves under a tank-in-series transport model along flow pathlines. These two approaches are complementary: the PINN targets richer hemodynamic inference, whereas IDT targets rapid residence-time mapping. The manuscript is organized as follows. Section 2 describes the dataset, the CFD ground-truth simulations, the PINN and IDT implementations, and the validation procedure. Section 3 presents flow visualizations and quantitative comparisons of both methods against CFD ground truth. Section 4 discusses the strengths and limitations of the proposed approaches and outlines potential extensions and clinical applications.

## 2 Materials and Methods

### 2.1 Image acquisition and segmentation

We retrospectively examined N = 6 patients from a previous study [20]. Each patient’s left atrial (LA) anatomy was segmented from time-resolved, three-dimensional computed tomography (i.e., 4D-CT) imaging following the standard protocols at each participating location (National Institute of Health, Bethesda, MD and University of California San Diego, La Jolla, CA). DICOM files, with spatial resolution between 0.32 and 0.48 mm in the *x* − *y* axial plane and 0.5– 1 mm in the *z*−direction, were generated by reconstructing the images following the CT scanner manufacturers’ standard algorithms. Time-resolved images were acquired at regular intervals throughout the cardiac cycle, corresponding to 5%–10% of the R–R interval. Three of the patients did not have left atrial appendage thrombus (LAAT) or a history of transient ischemic attacks (TIAs), and had normal LA function. The other three had impaired LA global function and two of them had an LAAT at the time of imaging while the other one had a history of TIAs. These patients are hereafter referred to as LAAT/TIA-negative and LAAT/TIA-positive, respectively. In the LAAT/TIA-positive patients thrombi were digitally removed prior to generating meshes for CFD simulations.

Each patient’s LA, left ventricle (LV), pulmonary vein (PVs) inlets, mitral valve (MV) outlet, and LAA were identified and segmented. These data were used to create inlets, outlets, and immersed-boundary Lagrangian meshes of the LA for the CFD simulations. LV and LA volumes were computed to determine the flow rates through the PV inlets and MV outlet as described below. Segmentation and meshing were performed using ITK-SNAP [43] and custom MATLAB scripts similar to our previous works [20, 28, 29]. Table 1 summarizes the anatomical and functional features parameters of the patients studied in our work and Fig. 1 shows the final segmented left atrial geometries from 4D CT images used for CFD simulations, PINN, and IDT framework.

**Table 1:** Anatomical and functional parameters of the cases reported in this study. *D*_MV_ is the mitral valve diameter 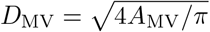 used as characteristic length, where *A*_MV_ is the area of the MV. *U*_MV_ is the peak velocity during the E-wave 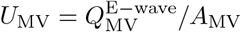 used as characteristic velocity, where *Q*_MV_ is the flow rate through the MV during E-wave. Reynolds number at the mitral valve annulus is calculated using the characteristic scales and the blood viscosity i.e., *Re* = *D*_MV_*U*_MV_*/ν*.

| Case Number | 1 | 2 | 3 | 4 | 5 | 6 |
| --- | --- | --- | --- | --- | --- | --- |
| LAA thrombus (or history of TIA) | No | No | No | TIAs | Yes | Yes |
| Atrial fibrillation | No | No | No | Persistent | Persistent | Persistent |
| Sinus rhythm | Yes | Yes | Yes | No | No | No |
| $D_{MV}$ (cm) | 2.14 | 1.78 | 2.40 | 2.42 | 2.43 | 2.33 |
| $U_{MV}$ (cm/s) | 80.44 | 140.77 | 105.37 | 72.15 | 67.71 | 138.40 |
| Reynolds number ($Re$) | 4297 | 6266 | 6315 | 4361 | 4111 | 8058 |

**Figure 1:**
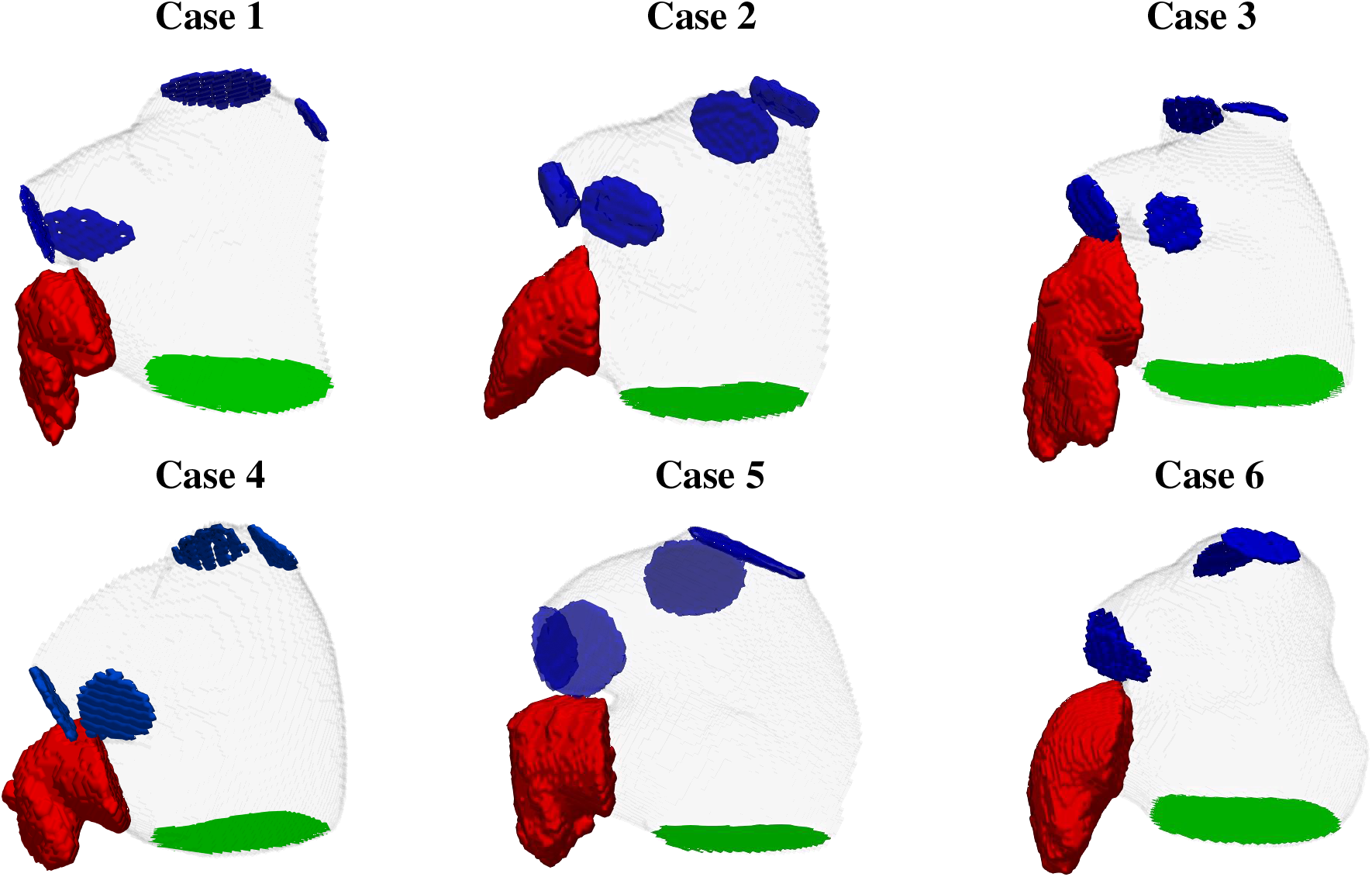
Patient-specific left atrial segmentations from computed tomography (CT) images depicting left atrium wall (in gray), left atrial appendage (in red), mitral valve plane (in green), and the pulmonary veins inlets (in blue). Top row shows the LAAT/TIA-negative cases (1-3) whereas the bottom row shows the LAAT/TIA-positive cases (4-6).

### 2.2 Computational fluid dynamics analysis

We used patient-specific CFD simulations to generate reference data for training and testing the PINN models and the IDT framework. Specifically, blood flow, contrast transport, and residence time within the LA were computed using TUCAN [20, 28, 29], an in-house solver for the partial differential equations (PDEs) governing mass conservation, momentum balance (Navier–Stokes), transport of contrast concentration (*c*), and residence time (*T*_*R*_),

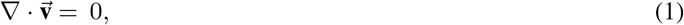

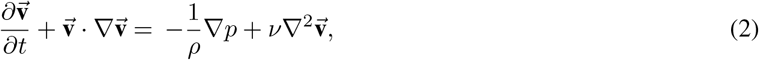

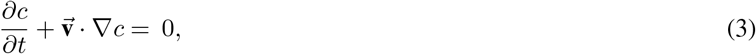

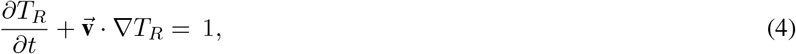

where 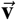 and *p* are the velocity and pressure fields. Blood was modeled as a Newtonian fluid of constant density *ρ* and kinematic viscosity *ν* = 0.04 cm^2^/s. The cardiac cycle was set to *t*_*c*_ = 1*s* in all cases. The contrast agent was modeled as a passive scalar, assuming it was too diluted to significantly affect blood flow via changes in density or viscosity. Residence time was computed by solving a transport equation with unit forcing, as previously described [44].

The simulations were initialized with 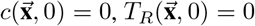 together with fully converged 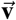 and *p* fields obtained after running our patient-specific CFD solver for *t* ≥ 20*t*_*c*_ in previous studies [20, 28, 29]. Starting from these conditions, both the Navier-Stokes and transport equations Eqs. (2)–(4) were integrated for an additional 60 *t*_*c*_ to simulate contrast bolus injection.

The bolus was prescribed through time-dependent contrast concentration profiles at the PV inlets. These profiles were derived from a gamma-variate model fitted to contrast-enhanced CT data from a separate patient, distinct from the six cases retrospectively analyzed here [37]. The fitted inlet profiles for the four PVs are shown in Fig. 2A. At the PV inlets, the residence time was set to zero and inflow velocities were set to 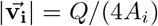 (i.e., even split), where *A*_*i*_ represents each PV’s cross-sectional area and *Q* is the total flow rate through the PVs, calculated from the patient-specific LA and LV volumes obtained by 4D CT. The motion of the LA wall obtained from the patient-specific 4D CT images was imposed via no-slip boundary conditions on 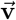. This no-slip condition included the mitral valve (MV) when it was closed. No particular spatial or temporal profile was considered for transmitral flow when the MV was open.

**Figure 2:**
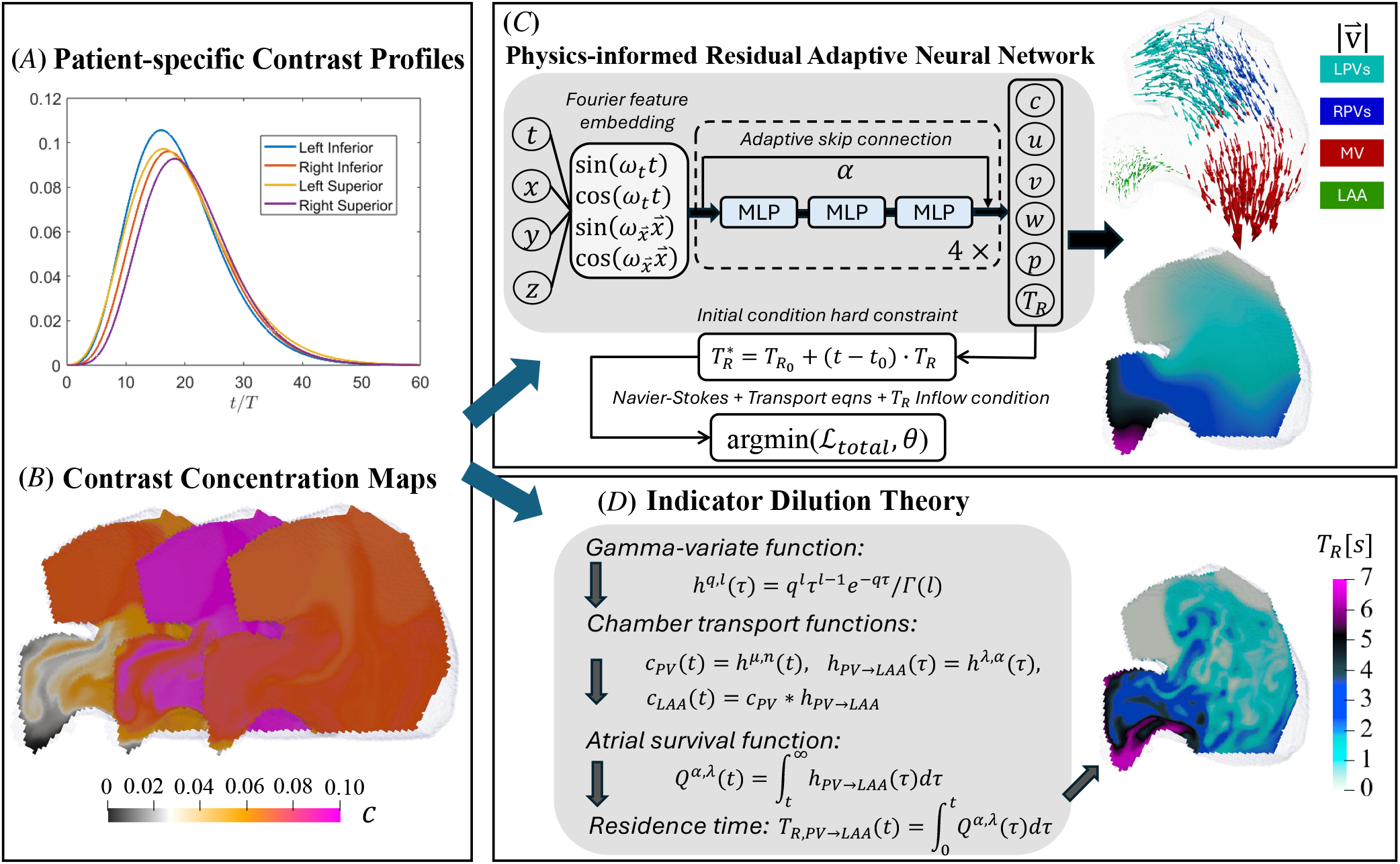
Overview of the study. *(A)* Patient-specific pulmonary vein (PV) contrast inflow profiles from 4D contrast CT are modeled with gamma-variate fits. *(B)* Contrast concentration maps obtained from CFD simulations. *(C)* Physics-informed residual adaptive neural network (PirateNet) framework: spatiotemporal inputs are encoded with Fourier features and passed through four residual blocks to infer contrast concentration, velocity, pressure, and residence time. Each block consists of three multilayer perceptrons (MLPs) and an adaptive skip connection step with a trainable parameter *α* initialized at 0. Training minimizes a composite loss including data, residence time inflow conditions, Navier-Stokes, and transport equations. *(D)* Indicator dilution theory (IDT) framework: chamber transport is modeled through an impulse response inferred from the PV inflow profiles and simulated contrast maps. Both inflow and response curves are parameterized with gamma-variate models, and residence time is obtained from the closed-form integral of the corresponding survival function (a hypergeometric function).

The patient-specific LA geometry was embedded in a cubic computational box of side length *L* = 13 cm. At the outer boundaries of this box, free-slip boundary conditions were imposed on 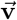, homogeneous Neumann boundary conditions were imposed on *c*, and homogeneous Dirichlet boundary conditions were imposed on residence time. Our formulation does not require boundary conditions for the pressure other than fixing its value at one point of the computational mesh. This value is immaterial since the pressure is undetermined by a constant in an incompressible flow.

Boundary conditions at the LA wall, mitral valve (MV), and pulmonary vein (PV) inlets were enforced with an immersed boundary method (IBM) [45], implemented as volumetric forcing terms in the corresponding evolution equations. At the LA wall, and at the MV plane when the valve was closed, the IBM forcing enforced no-slip conditions on 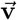. When the MV opened, this forcing was removed, yielding an idealized instantaneous opening/closing model. At each PV inlet, the prescribed inflow was imposed through a cylindrical buffer region composed of 12 cross-sectional layers extending upstream along the local inlet normal. A smoothed forcing term applied over this buffer region drove the flow toward the target inlet velocity, following [20].

We integrated the evolution equations (2)–(4) in time using a low-storage, three-step, semi-implicit Runge-Kutta. The time step was set constant to the Courant-Friedrichs-Lewy (CFL) number below 0.2. Continuity was enforced by splitting the temporal integration of the Navier-Stokes equations using a fractional step method with pressure correction. The spatial derivatives in the Navier-Stokes equations were approximated using second-order finite differences on a uniform Cartesian staggered grid. A third-order WENO scheme was used for the spatial discretization of convective terms in the transport equations (3)–(4) to mitigate spurious oscillations near sharp gradients [46]. The spatial resolution was Δ*x* = 0.050 cm, matching that used in previous simulations [20, 28, 29]. To reduce the computational cost of PINN training, the CFD results were subsequently downsampled to Δ*x* = 0.090 cm and used as the reference dataset for PINN training and validation.

### 2.3 Physics-informed neural network

#### 2.3.1 Architecture

We built a physics-informed neural network (PINN) with PirateNet [47] as the underlying architecture to infer flow and residence time from the 4D contrast dynamics obtained using CFD. PirateNet is a residual architecture with adaptive residual connections that improves stability and mitigates the deep multilayer perceptrons (MLP) derivatives’ initialization difficulties in training deep PINNs. Through its trainable residual connections, PirateNet initially learns toward a stable linear combination representation and progressively recovers richer nonlinear expressiveness during training. In our study, the PINN approximates the mapping function 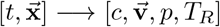, and the underlying PirateNet is composed of 4 residual blocks, each containing 3 MLPs with 200 neurons per layer and the nonlinear *swish* activation function [48]. This deep and dense configuration was chosen because of the complexity of the geometries and the underlying flow dynamics. The neural network parameters (i.e., weights) were initialized using Xavier’s scheme [49], and weight normalization [50] was employed to accelerate training.

We employed Fourier feature embeddings to enforce exact temporal periodic condition and to enrich input spatial features proposed in previous literature to circumvent spectral bias in the PINN [51]. Specifically, the spatiotemporal contrast inputs were passed through a structured Fourier embedding layer comprising sine and cosine functions,

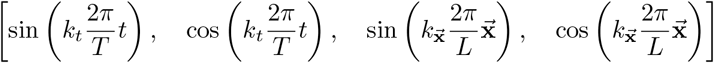

where 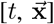 are spatiotemporal inputs, *T* is the period of the cardiac cycle, and *L* is the domain size. *k*_*t*_ and 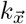 are array of mode indices (i.e., *k*_*t*_ = [1, 2, 3, …, *M*_*t*_] and 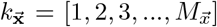. We chose the number of modes in the Fourier layer *M*_*t*_ = 2 and 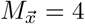. This configuration was selected empirically to balance computational cost and high frequency Fourier feature expressibility in the MLP [51]. Higher Fourier modes were also evaluated for 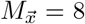 and 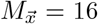 to encode finer spatial features but they did not lead to improved accuracy or performance.

To impose residence time initial condition in the PINN, we chose to impose it as a hard constraint [52] by reparameterizing the network output as as,

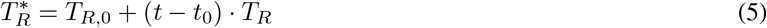

where *T*_*R*_ denotes the raw output of the MLP, *t*_0_ is the initial time, and *T*_*R*,0_ is the desired residence time map at *t*_0_; here, *T*_*R*,0_ = 0 everywhere. The reparameterized variable 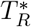 was then used as the residence time field in Eq. 4. This construction enforces the initial condition exactly, rather than penalizing deviations from it through an additional loss term. For simplicity, we continue to denote the residence time field as *T*_*R*_ throughout the manuscript.

The model was trained by minimizin the composite loss function

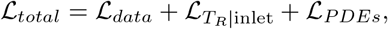

which penalizes mismatches between the predicted contrast concentration *c* and the training data (ℒ_*data*_), departures from zero residence time at the PV inlets 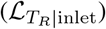, and residuals of the PDEs governing blood flow and transport of *c* and *T*_*R*_ (ℒ_*PDEs*_).

At each training iteration, the optimizer sampled the CFD-derived 4D contrast dataset *c*_data_ to evaluate the data-fidelity term,

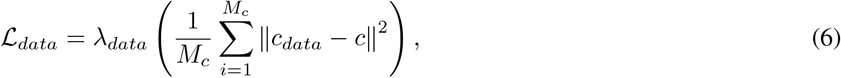

where *λ*_*data*_ is a weight coefficient and *c* is the MLP contrast output. The training batch comprised *M*_*c*_ randomly drawn points from the CFD mesh across *N*_*Cca*_ frames.

The loss term enforcing the inlet condition on residence time was defined as

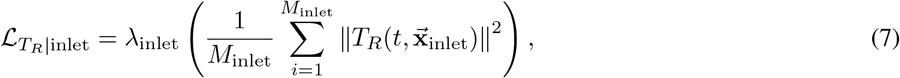

where *λ*_inlet_ is the corresponding weight. For this term, all available points at the four PV inlets were sampled across all frames.

The loss terms corresponding to the residuals of the governing equations were defined as

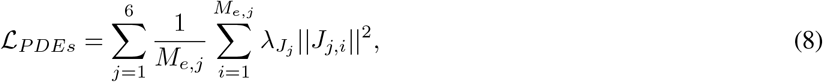

where 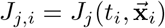 represents the residuals of the *j*-th PDE,

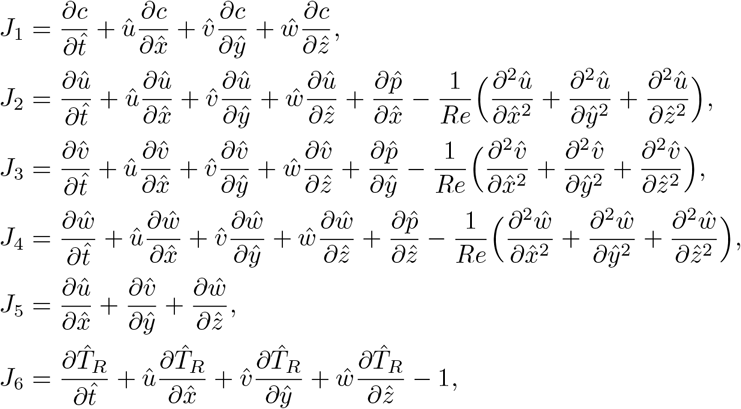

obtained at *i*-th the sampling point, 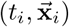. The partial derivatives were computed using automatic differentiation [53]. The residuals were computed in non-dimensional form to improve training convergence [35, 54]. Specifically, we used the change of variables,

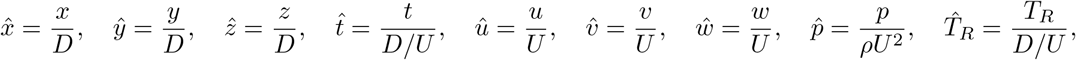

where *U* is the peak mitral-valve velocity during rapid LV filling (E-wave), *D* is the mitral-valve diameter, *ρ* is the blood density, and *Re* = *DU/ν* is the Reynolds number. These characteristic scales are listed in Table 1. In addition, all variables, including the contrast training data *c*, were standardized to zero mean and unit variance to improve training robustness and reduce vanishing-gradient issues [49]. After training, the inferred fields were transformed back to physical units. The same nondimensionalization and normalization procedures were applied to all loss terms. Equation residuals were evaluated at *M*_*e*_ collocation points, randomly resampled across space and time from the available *N*_*Cca*_ and *N*_*eqns*_ frames at each training iteration.

#### 2.3.2 Patient-specific training and validation

Contrast released at the PV inlets required approximately 10 cardiac cycles to fill a substantial portion of the LA. Accordingly, frames with *t* ≤ 10 *t*_*c*_ were excluded, and for each patient-specific case the loss terms were sampled from cycles 10 through 25 using *N*_*Cca*_ = *N*_*eqns*_ = 300 frames. For each iteration, we used batches of size *M*_*c*_ = *M*_*e*_ = 10^4^ randomly sampled points, and the composite loss function was minimized with the *Adam* optimizer [55] Training was initialized with a learning rate of 10^−3^ for 10^5^ iterations, followed by a fine-tune phase using a learning rate of 10^−4^ for additional 5 *×* 10^4^ iterations as illustrated for a representative case in Fig. SI1. The weighting coefficients associated with all loss terms were set equal to one (*λ*_*data*_ = 1, etc.) in all cases. The PINN framework was implemented in TensorFlow [56], building on our previous work [57], and training was performed on an NVIDIA A100 GPU with 80 GB of memory. Total runtime ranged from 10 to 15 hours per case. All other processing and visualization steps were carried out using in-house MATLAB and ParaView scripts.

Correlation coefficients *r* were computed for the velocity fields and residence-time maps, and the corresponding statistical distributions of kinetic energy density, *KE* = (*u*^2^ + *v*^2^ + *w*^2^)/2, and residence time were also compared. In addition, velocity-field discrepancies between the contrast-trained PINN and the CFD reference were quantified using the normalized magnitude (*ϵ*_*M*_) and orientation (*ϵ*_*O*_) errors:

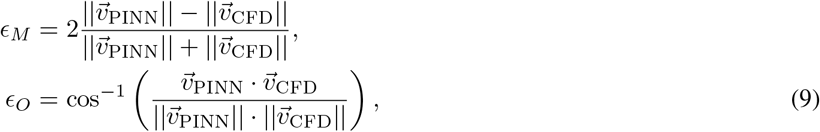

where 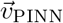 and 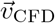 denote the velocity fields predicted by the contrast-trained PINN and computed by CFD, respectively. The orientation error measures the angle between 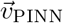 and 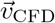. Positive values of *ϵ*_*M*_ indicate larger velocity magnitudes in the PINN than in CFD, and *vice versa*.

### 2.4 Indicator dilution theory

#### 2.4.1 Chamber response and residence time derivations

Indicator dilution theory (IDT) [58] models tracer transport between an injection site **x**_*i*_ and a downstream location **x** as a statistically stationary linear process, fully characterized by a transport function 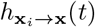 [59]. Accordingly, the tracer concentration at **x** is given by the convolution of the injection concentration profile 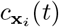 with the transport function,

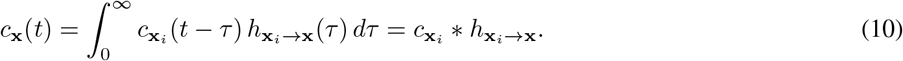

The transport function represents the system’s impulse response to a Dirac delta bolus injection. It has been commonly represented by a gamma-variate function following a tank-in-series model [60],

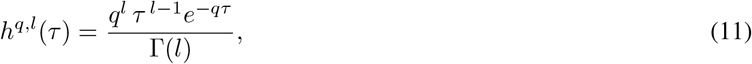

where *l* and *q* denote the shape and rate parameters, respectively. These parameters are often interpreted phenomenologically as reflecting the effective volume and flow rate of the compartment, and Γ(*·*) denotes the gamma function. Assuming a bolus injection that delivers a finite dose over a time interval that is short relative to the transport dynamics, the injection profile 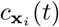 may be idealized as a Dirac delta function. In this case, the observed concentration *c*_**x**_(*t*) coincides with the transport function itself. Accordingly, empirical concentration–time curves are often fitted with gamma-variate models to estimate the underlying transport function [42].

The associated survival function

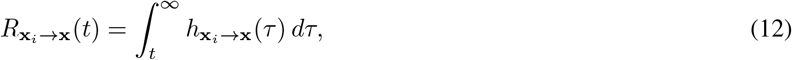

quantifies the fraction of tracer remaining within the compartment bounded by **x**_*i*_ and **x** at time *t*. The corresponding residence time is defined as

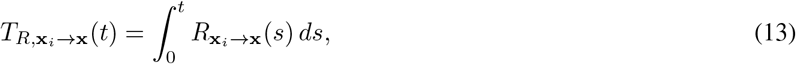

and its limit when *t* → ∞ is the expectation of the transport function,

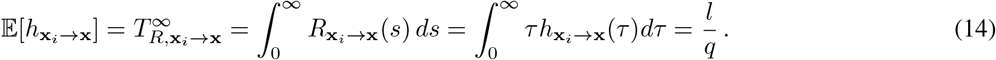

The expectation and variance of the transport function are additive, i.e.,

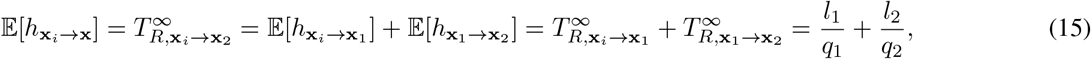

and

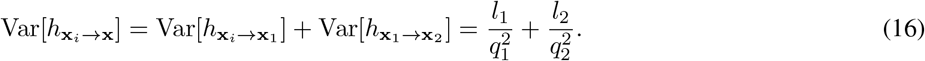

In contrast, the convolution of two gamma-variate transport functions only yields another gamma-variate function when they have the same rate parameter, in which case the shape parameter becomes additive, i.e.,

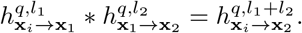

We seek to apply IDT to model contrast dynamics within the LA voxel by voxel, using the contrast profile measured at the pulmonary veins (PVs) as the inlet reference to determine the transport function *h*_PV→LAA_(*τ*). This approach introduces a critical challenge: the PV concentration profile is not a Dirac delta input, and *h*_PV→LAA_ may not share the same rate parameter as the upstream transport function 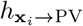, which reflects the branched transport from the bolus injection site, typically a peripheral vein in the arm, to the PVs. Therefore, modeling *c*_LAA_(*t*) as a gamma-variate function would not directly yield *h*_PV→LAA_(*τ*) nor the corresponding residence time 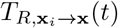. Accordinly, we assume that both the PV inlet contrast profile and the PV → LAA transport function follow gamma-variate forms,

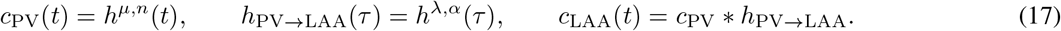

To determine the parameters *α* and *λ* of the atrial transport function, we employ an integral-moment–based method that enables fast and robust estimation without explicit fitting of temporal concentration curves. Leveraging the additive properties of the mean and variance (Eqs. 15–16), we write

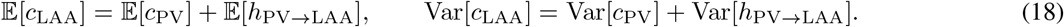

Assuming that both *c*_PV_(*t*) and *c*_LAA_(*t*) can be measured, Eqs. (17)–(18) allow the transport parameters to be computed as

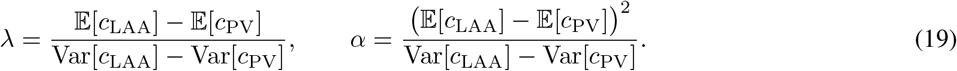

To derive the LAA residence time, we define the atrial survival function

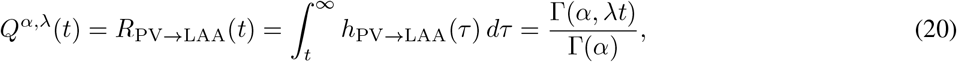

where Γ(*α, λt*) denotes the upper incomplete gamma function, which is fully determined by the transport parameters *α* and *λ*.

The residence time is then obtained by integrating the survival function,

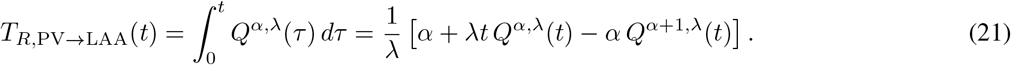

This closed-form expression satisfies the initial condition *T*_*R*,PV→LAA_(0) = 0, since *Q*^*α,λ*^(0) = 1 provided that *h*_PV→LAA_ is a properly normalized probability density function. In the limit *t* → ∞, the last two terms in Eq. (21) cancel each other out, yielding the expected result

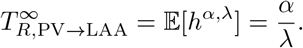

An alternate working solution to the challenge created by the PV concentrations not being dirac Deltas is to use gamma-variate fits to estimate the transport functions 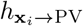 and 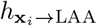, from which 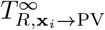 and 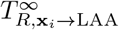 can be inferred. The residence time between the PVs and the LAA is then obtained as 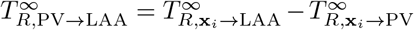 [37]. However, this approach defines the origin of residence time at the instant of bolus injection rather than at arrival at the PVs, and is therefore valid only in the limit *t* → ∞. This estimate can overestimate residence time at finite times when comparing to CFD data from finite-time simulations.

#### 2.4.2 Assumptions and rationale for applying IDT in periodic 3D atrial transport

Our IDT formulation approximates LA blood transport by an effective linear tank-in-series process when analyzed at a fixed cardiac phase. Its voxel-wise application to 3D LA flow departs from traditional applications to 0D representations of the circulatory system, for which the assumptions of linearity and stationarity have been empirically validated [61]. Here, we reason that the 0D approximation can be extended to 3D under two assumptions: (i) the transport process is stationary in the mean across cycles, and (ii) contrast behaves as a passive tracer with negligible molecular diffusion (high Péclet number). Neglecting diffusion is consistent with the low diffusivity of iodine contrast agents in blood, and is important because it allows residence time to be inferred from temporal moments of advective transport alone, without introducing variables that could confound the identification and interpretation of the transport parameters (*q, l*).

Under these assumptions, we write *t* = *t*\**T* +*τ* with *τ* ∈ [0, *T*), where *t*\* is a slow time variable (cycle count)and *τ* is within-cycle time. Fixing a phase *τ*_0_ and sampling once per cycle at that phase defines the phase-locked field 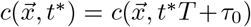. By flow periodicity, 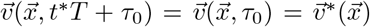, so 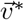 is independent of *t*\*. The phase-locked contrast dynamics across cycles is governed by

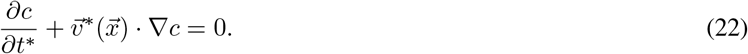

To motivate the tank-in-series model in this setting, consider transport restricted to a phase-locked pathway coordinate *s* with local advective speed *u*(*s*), so that

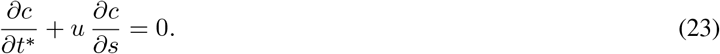

Discretizing the advective term in Eq. (23) with a first-order upwind scheme on a grid *s*_*j*_ = *j* Δ*s* yields the semi-discrete system

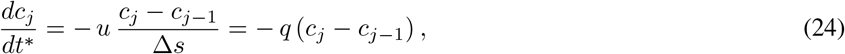

i.e., a cascade of *l* first-order relaxation (mixing) elements in series with per-stage rate *q* = *u/*Δ*s* (or locally *q*_*j*_ = *u*(*s*_*j*_)/Δ*s*), which motivates the gamma-variate (tank-in-series) impulse response used in §2.4.1. The associated numerical diffusivity of the upwind discretization, *D*_num_ = *u* Δ*s/*2 = *u*^2^/(2*q*), quantifies the dissipative coarse-graining implicit in the upwind/tank analogy rather than physical dispersion. Moreover, it produces no transverse mixing between trajectories consistent with the high Péclet number approximation, since it acts only along the pathway coordinate *s*. Finally, for a pathway section length *L*_*s*_ the *l*-stage tank-in-series response has a transit time given by 𝔼 [*h*] = *l/q* = *L*_*s*_*/u*, which is independent of Δ*s* and therefore independent of *D*_num_.

#### 2.4.3 Voxelwise IDT Implementation

To obtain voxel-wise residence times, we apply the tank-in-series model along the flow pathlines, effectively extending it to 3D unsteady flow. Accordingly, we define

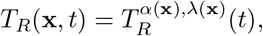

where *α*(**x**) and *λ*(**x**) are obtained by applying the integral-moment relations in Eq. (19) to each voxel’s contrast profile *c*(**x**, *t*). For each patient’s PV, we fit the inlet contrast time curve (Fig. 2A) with a gamma-variate model using nonlinear least squares to estimate the shape and rate parameters (*n, µ*). Because each left atrium has multiple (typically four) PV inlets, we used the mean values of *n* and *µ* across PVs as the inlet reference for the downstream moment calculations.

The full CFD contrast dataset spanning 60 cardiac cycles, ensuring that both the wash-in and wash-out phases were included. To compute voxelwise mean and variance, we grouped the contrast concentration fields by cardiac phase (i.e., the ℓ-th frame within each cardiac cycle) and stacked the resulting time series. This procedure emulates clinical low-dose multi-cycle 4D contrast CT acquisitions, where only one 3D volume is reconstructed at a given cardiac phase per heartbeat (or every ~ *k* heartbeats), due to temporal-resolution and radiation-dose constraints [37]. For each voxel and phase, the stacked concentration curve was normalized by its temporal integral and then integrated in time to compute the mean *m*_*c*_ and variance *v*_*c*_. These moments, together with the inlet reference moments, were used to compute the voxelwise transport parameters *α* and *λ* via Eq. (19). Finally, the residence time evolution at each voxel was obtained from the closed-form expression in Eq. (21). All fitting and computations were performed using in-house MATLAB scripts; additional details are provided in Algorithm 1.

##### Algorithm 1

Residence time computation via indicator dilution theory

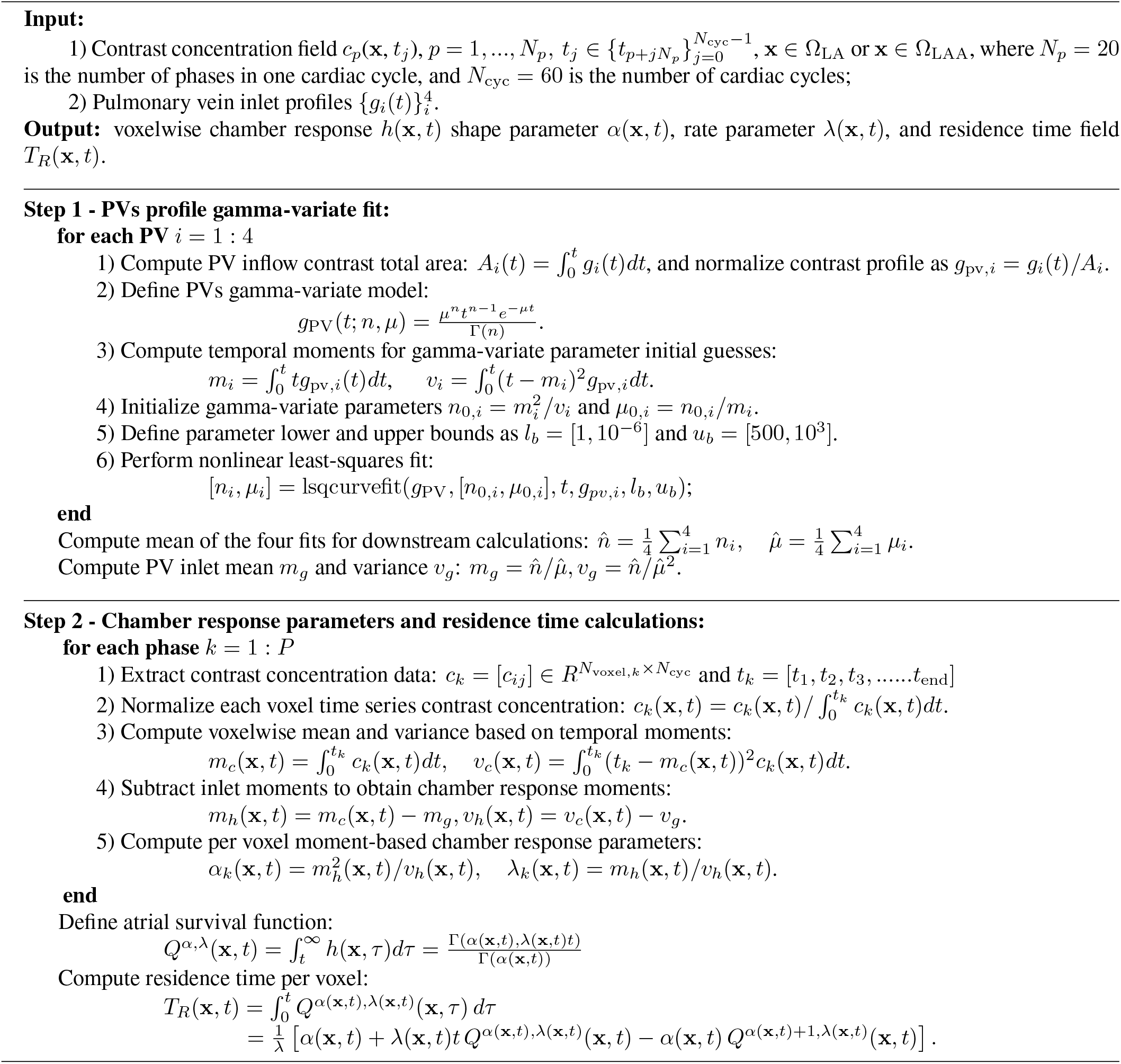

## 3 Results

This section presents flow visualizations and quantitative validation of the PINN and IDT methods, using CFD-derived contrast concentration fields for training, and residence time and flow fields for validation. For brevity, we highlight flow visualizations from two representative cases: one LAAT/TIA-negative patient (case 3) and one LAAT/TIA-positive patient (case 5), while reporting quantitative statistics across all six cases. Additional visualizations for the remaining cases are provided in the Supporting Information. Overall, the PINN accurately reconstructs the dominant LA flow structures across the cardiac cycle and yields residence time maps *T*_*R*_ that show reasonable agreement with CFD, with a tendency to underestimate the largest *T*_*R*_ values, and requiring hours of patient-specific training. IDT does not recover velocity fields, but infers residence time maps from contrast curves within seconds with excellent agreement with CFD. Despite these differences, both methods discriminate between LAAT/TIA-negative and LAAT/TIA-positive cases based on the inferred LAA *T*_*R*_.

### 3.1 Spatiotemporal Comparison of *T*_*R*_ Fields: CFD vs. PINN and IDT

Figure 3 shows snapshots of the contrast concentration and residence time fields, 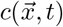 and 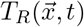, from the CFD simulations for representative cases 3 (LAAT/TIA-negative) and 5 (LAAT/TIA-positive). The corresponding PINN-inferred fields are also shown, together with IDT-derived 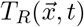. Contrast concentration snapshots are shown at *t*_*c*_*/T* = 12, 18, and 25, whereas residence time snapshots are shown at 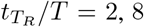, and 15. Here *t*_*c*_ is referenced to the contrast-training window (cycles 10–25), and 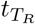 is referenced to the start of residence time integration. Each snapshot is selected at the same phase of the cardiac cycle (early atrial diastole) but spanning different cycles. Analogous maps for the remaining four cases are provided in the Supporting Information (Figs. SI5–SI7). Because the contrast data-misfit term directly constrains 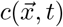 during training, the PINN reproduces the contrast concentration with minimal deviation from the CFD reference across cases and time points.

**Figure 3:**
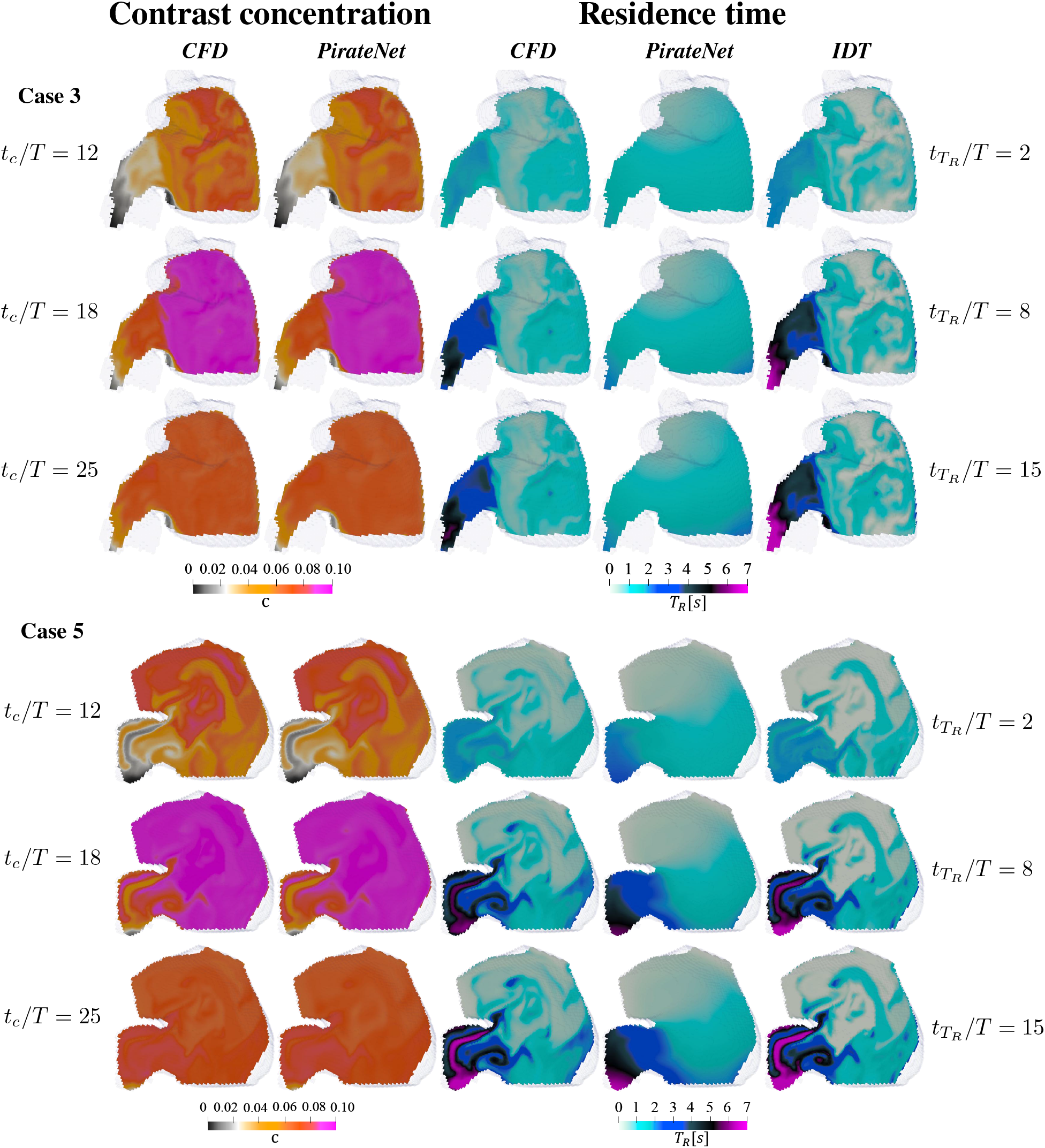
Contrast concentration (*c*) and residence time (*T*_*R*_) maps comparison of case 3 (top) and case 5 (bottom) between CFD, PirateNet, and IDT. Three 2D sections of the left atrium are shown at selected time instants. Contrast concentration snapshots are shown at *t*_*c*_*/T* = 12, 18, and 25, whereas residence time snapshots are shown at 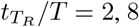, and 15. Here *t*_*c*_ is referenced to the contrast-training window (cycles 10–25), and 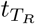 is referenced to the start of residence time integration. Contrast concentration is dimensionless, whereas residence time is given in seconds.

In the LAAT/TIA-negative case (top panels of Fig. 3), during the early stages of bolus transport (*t*_*c*_ = 12*T*) contrast enters the LA through the PVs but does not yet penetrate deeply into the LAA, so 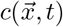 remains low and largely confined to the LA body. At this time, the PINN recovered the overall spatial organization of 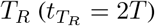, but it smoothed fine-scale features and tended to underestimate *T*_*R*_ within the LAA. By comparison, IDT yielded a more detailed *T*_*R*_ map that more closely matched the CFD reference.

As the bolus continues to enter through the 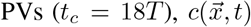 increases throughout the LA body and begins to rise within the LAA, with a delayed wash-in relative to the LA. This lag is consistent with longer residence times in the appendage, as reflected by the CFD results. At this stage 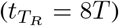, the PINN reproduced the overall 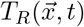 distribution but underestimated *T*_*R*_ within the LAA. Notably, the IDT reconstruction resolved the sharp *T*_*R*_ gradient at the LAA ostium, in close agreement with the CFD reference, while tending to overestimate *T*_*R*_ in distal regions. At later cycles 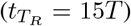, the PINN still underestimated *T*_*R*_ in case 3, particularly in high-residence-time regions of the LAA. On the other hand, IDT remained closer to the CFD reference and continued to capture the ostial *T*_*R*_ gradient, although it slightly overestimated distal *T*_*R*_ values.

The performance of both inference methods was more favorable in the LAAT/TIA-positive case (bottom panels of Fig. 3). Across all reported time points during bolus wash-in and wash-out, both PINN and IDT yielded 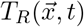 fields in good visual agreement with the CFD reference, although the PINN again appeared smoother and missed some fine-scale structure. The agreement between IDT-derived and CFD-derived residence time was remarkable.

### 3.2 Statistical Validation of Inferred LAA *T*_*R*_

Given the good agreement of PINN- and IDT-derived *T*_*R*_ patterns in the LA body, we focus our quantitative evaluation on the LAA, the predominant site of thrombus formation and the region most sensitive to prolonged residence time. We therefore estimate the joint probability density functions relating inferred and CFD residence times within the LAA, i.e., *p*(*T*_*R*,CFD_, *T*_*R*,PirateNet_) and *p*(*T*_*R*,CFD_, *T*_*R*,IDT_), and the corresponding correlation coefficients *R*_CFD,PirateNet_ and *R*_CFD,IDT_. These statistics are calculated by pooling all LAA voxels over 15 cardiac cycles. Fig. 4 displays contours of *p*(*T*_*R*,CFD_, *T*_*R*,PirateNet_) and *p*(*T*_*R*,CFD_, *T*_*R*,IDT_) for the six patient-specific cases analyzed. The outermost PDF contour contains 99% of the sampled points and the innermost one contains 75%. In line with the map-level comparisons in Fig. 3, IDT exhibited near-uniform accuracy across the full range of *T*_*R*_ values, with deviations confined to an overestimation of the upper tail. The PINN, in turn, showed good agreement with CFD for low-to-intermediate residence times but progressively underestimated *T*_*R*_ as values increase, leading to a compression of the high-*T*_*R*_ range. This underestimation was more severe in the in the LAAT/TIA-negative cases (Fig. 3A–C), where the PINN rarely predicted a *T*_*R*_ value higher than 4 cycles. As a result, the PINN provided comparatively better residence-time reconstructions in the LAAT/TIA-positive cohort. Violin plots of the univariate distributions *p*(*T*_*R*_) from CFD, PINN, and IDT (Supplementary Fig. SI8) confirmed these trends. Likewise, the computed correlation coefficients (Table 2) indicated that IDT achieved higher correlations with CFD (*r* = 0.81–0.94) than the PINN (*r* = 0.51–0.82), and the PINN correlations were obtained highest in the LAAT/TIA-positive cases than in the LAAT/TIA-negative cases.

**Figure 4:**
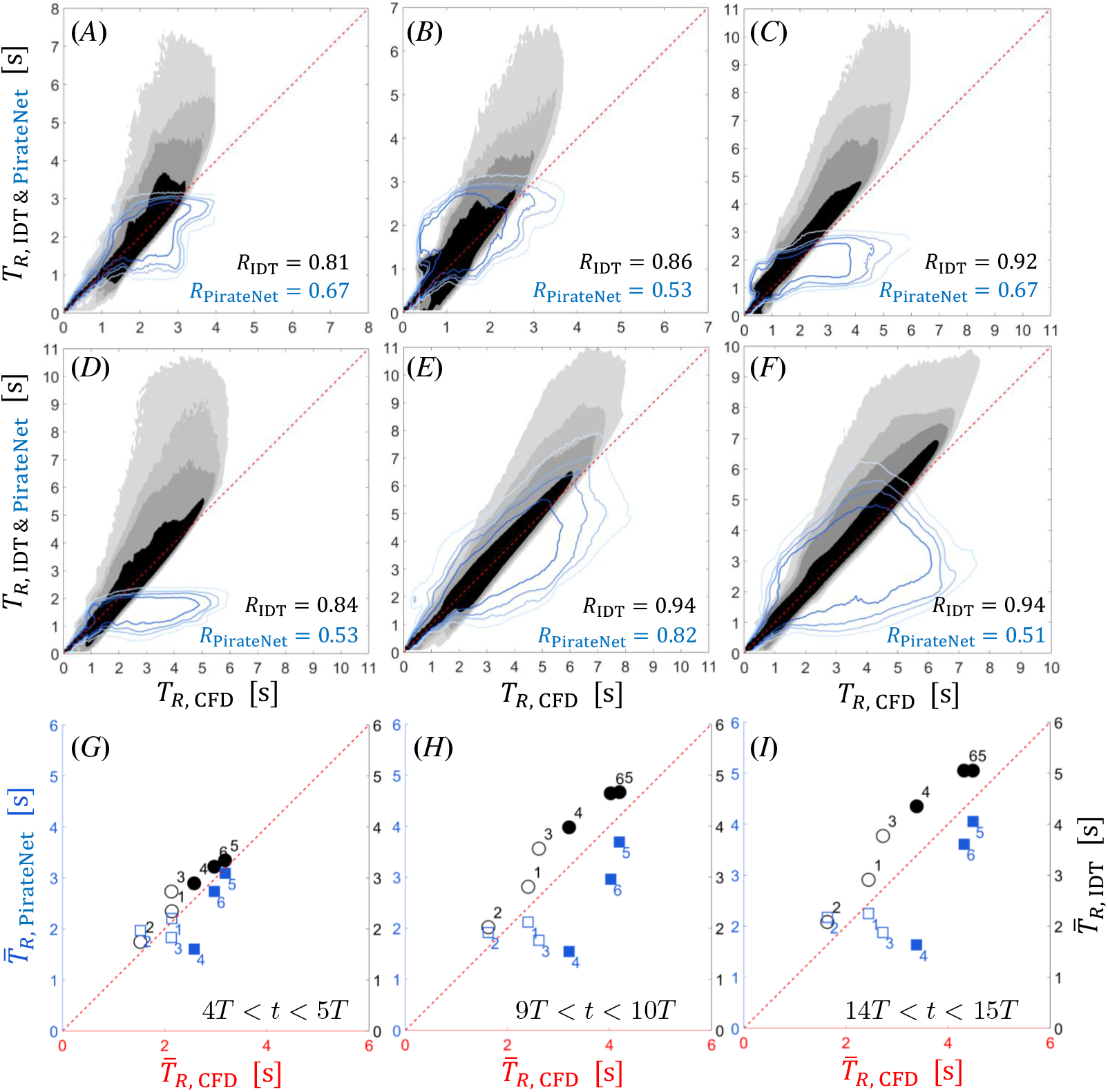
Comparison of LAA *T*_*R*_ by PirateNet and IDT with CFD ground-truth data. The top panels (*A*–*F*) display joint probability density functions of CFD residence times (*x*-axis) and IDT-inferred residence time (*y*-axis, filled black contours), *p*(*T*_*R*,CFD_, *T*_*R*,IDT_), or PirateNet-inferred residence time (*y*-axis, blue line contours), *p*(*T*_*R*,CFD_, *T*_*R*,PirateNet_). 15 cycles of residence time are pooled to display the joint pdfs. Panels (*A*) through (*F*) respectively represent cases 1 through 6. Each contour, from dark to light, contains 75%, 90%, 95%, and 99% of the data, respectively. The bottom right corner of each panel shows the correlation coefficients *R*_PirateNet_ and *R*_IDT_. The bottom panels (*G*–*I*) are scatter plots of mean LAA *T*_*R*_ from PirateNet and IDT vs. with mean *T*_*R*_ from CFD, averaged within the LAA over three different cardiac cycles: *G*), 4*T* ≤*t* ≤ 5*T*; *H*), 9*T* ≤ *t* ≤ 10*T*; *I*) 14*T* ≤ *t* ≤ 15*T*. Hollow and solid symbols respectively represent LAAT/TIA-negative and cases LAAT/TIA-positive cases.

**Table 2:** Global velocity comparison metrics between PINN and CFD for the six studied cases. PINN results are reported for both PirateNet and vanilla MLP architectures. *ϵ*_*M*_ represents the normalized magnitude discrepancy and *ϵ*_*O*_ is the global orientation error (in degrees). LAA residence time *T*_*R*_ correlations are shown for PINN vs. CFD and IDT vs. CFD.

| Case | $T_{R,LAA}$ correlation coefficient | | | Velocity errors | | | | | | | |
| --- | --- | --- | --- | --- | --- | --- | --- | --- | --- | --- | --- |
| | PINN vs. CFD | | IDT vs. CFD | $\epsilon_M^{LA}$ (-) | | $\epsilon_O^{LA}$ (°) | | $\epsilon_M^{LAA}$ (-) | | $\epsilon_O^{LAA}$ (°) | |
|  | PirateNet | MLP | – | PirateNet | MLP | PirateNet | MLP | PirateNet | MLP | PirateNet | MLP |
| 1 | 0.67 | 0.65 | 0.81 | 0.98 | 0.99 | 32.04 | 32.64 | 0.88 | 0.93 | 38.81 | 38.02 |
| 2 | 0.53 | 0.37 | 0.86 | 0.90 | 0.81 | 29.26 | 28.15 | 0.61 | 0.56 | 35.90 | 36.35 |
| 3 | 0.67 | 0.63 | 0.92 | 0.82 | 0.77 | 28.43 | 28.19 | 0.74 | 0.73 | 30.86 | 29.39 |
| 4 | 0.53 | 0.59 | 0.84 | 0.56 | 0.48 | 27.37 | 27.58 | 0.47 | 0.48 | 25.34 | 30.62 |
| 5 | 0.82 | 0.84 | 0.94 | 0.35 | 0.35 | 19.92 | 19.27 | 0.35 | 0.35 | 23.64 | 23.27 |
| 6 | 0.51 | 0.51 | 0.94 | 0.76 | 0.79 | 30.68 | 30.49 | 0.38 | 0.40 | 27.96 | 28.31 |

To assess whether the PINN and IDT methods can discriminate LAAT/TIA-positive from LAAT/TIA-negative cases despite their biases, we compared mean LAA residence times against CFD using scatter plots (Fig. 3G–I). These data confirmed the trends observed above: IDT tended to overestimate mean *T*_*R*_, whereas the PINN tended to underestimate it. Notably, case 3— used for the detailed flow and *T*_*R*_ visualizations—showed one of the strongest manifestations of these biases in panels G–I. Despite these offsets, the inference was favorable relative to the intrinsic inter-patient variability in LAA residence time. Importantly, both methods preserved the clinically relevant ordering across patients: PINN- and IDT-inferred mean LAA *T*_*R*_ values are higher in LAAT/TIA-positive cases than in LAAT/TIA-negative cases. This behavior was conserved across a different cardiac cycles as the bolus washes in and out the LA and LAA, with a trend for biases to increase as *T*_*R*_ increases (compare panels G–I in Fig. 3).

### 3.3 Temporal Dynamics of Inferred LAA *T*_*R*_

The temporal evolution of LAA residence time in CFD typically shows two regimes [20]. Starting from *T*_*R*_ = 0, residence time initially grows approximately linearly over a period set by the advective delay for tracer to reach the LAA from the PVs (see insets in Fig. 5). Once the LAA begins to exchange fluid significantly with the atrial body, *T*_*R*_ grows more slowly and is modulated by cardiac-cycle oscillations, consistent with dispersive mixing. Over longer times, the mean *T*_*R*_ approaches a quasi-steady plateau, whose level reflects the characteristic time required to renew the LAA blood pool. This behavior is illustrated in Fig. 5, which shows LAA volume-averaged residence time, 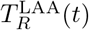 for each case over 15 cardiac cycles. Shown in this way, the data tests whether the PINN and IDT frameworks trained on contrast can reproduce the temporal dynamics of 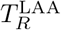.

**Figure 5:**
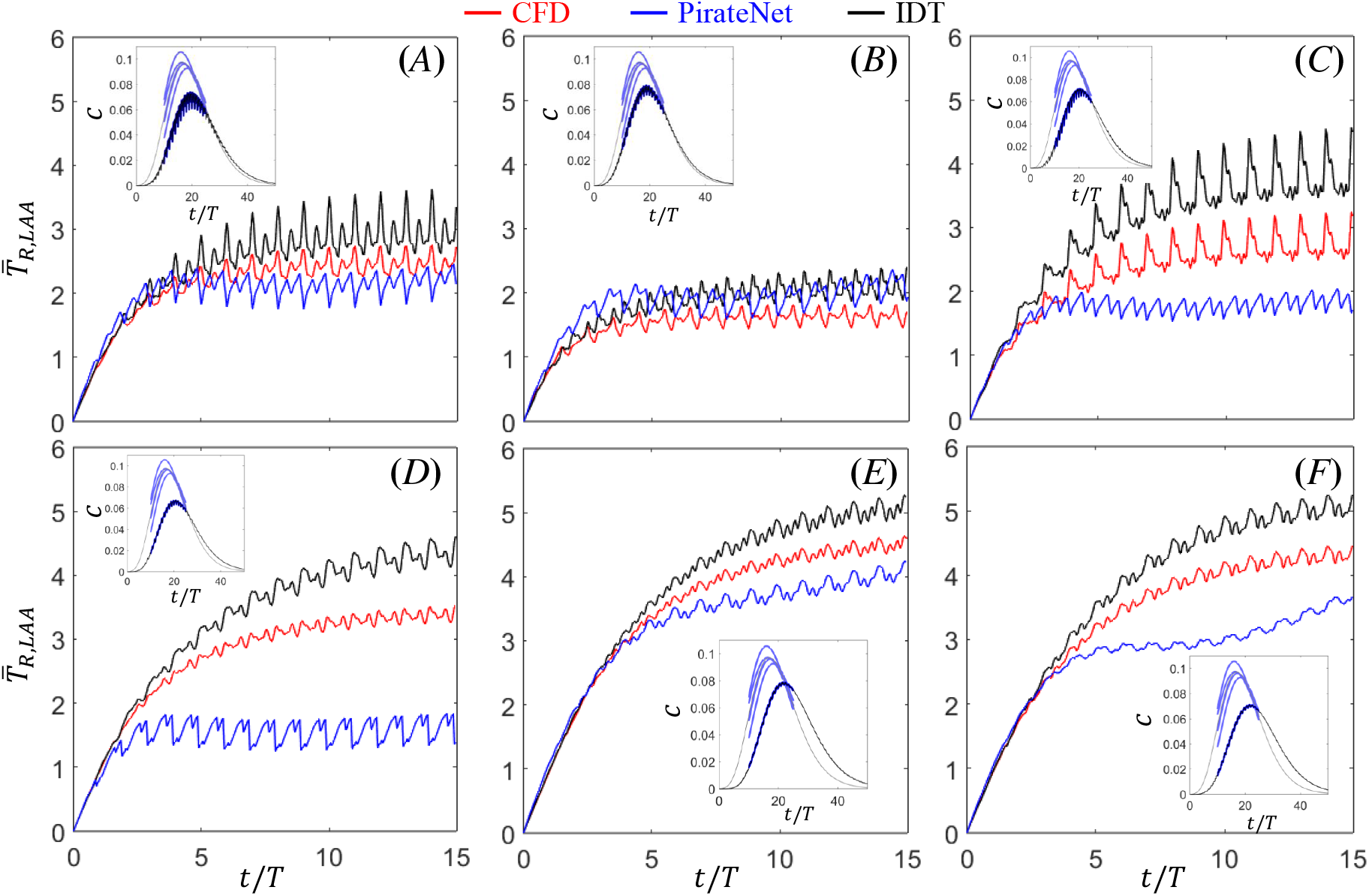
Comparison of volume-averaged LAA residence time *T*_*R*_ inferred by the PirateNet (blue) and IDT (black) against CFD ground truth (red). Panels (*A*)–(*F*) correspond to cases 1–6. Insets show representative training contrast concentration time curves at inlet and chamber locations (lighter/darker shades indicate PVs/LAA). For the PINN, the LAA volume-averaged contrast and the cross-section–averaged contrast at each PV inlet are shown. For IDT, the LAA volume-averaged contrast and the cross-section–averaged contrast pooled across the four PV inlets are shown. In each inset, only the time interval used to train the corresponding method is shown.

Overall, both PINN- and IDT-inferred 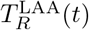 captured the initial linear-growth regime and its duration. In the subsequent dispersive regime, both methods also reproduced the pulsatile modulation of 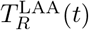, including the correct cardiac frequency and the larger oscillation amplitude in LAAT/TIA-negative than in LAAT/TIA-positive cases. However, the two methods differed in the slow-time evolution of the mean 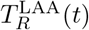 in the dispersive regime. The PINN tended to flatten earlier, yielding a lower long-time envelope, whereas IDT relaxed more gradually and remained higher over the same window. These differences translated into a net underestimation of LAA *T*_*R*_ by the PINN and a net overestimation by IDT, consistent with the biases identified in the previous section.

### 3.4 Left Atrial Flow Reconstruction Using PINN

We next assessed the ability of the PINN and IDT frameworks to resolve flow within the LA and LAA. The PINN directly inferred flow velocities while the flow rate parameter *λ* inferred by IDT method could be interpreted as a measure of local flow velocity magnitude. However, spatial maps of voxel-wise inferred rate parameter, *λ*(**x**, *t*), not shown, were not informative of flow patterns. Therefore, this section focuses on PINN-inferred velocities.

Figure 6 compares CFD and PINN instantaneous LA velocity fields at three representative phases of the cardiac cycle (atrial diastole, early LV filling, and atrial systole) for case 3 case 5. Vector fields from both sources as well as relative differences in magnitude (*ϵ*_*M*_) and orientation (*ϵ*_*O*_), defined in equations (9), are presented. The accompanying pulmonary vein (PV) and mitral valve (MV) flow rate waveforms, and LA and LAA volume traces, provide hemodynamic context for the selected time instants.

**Figure 6:**
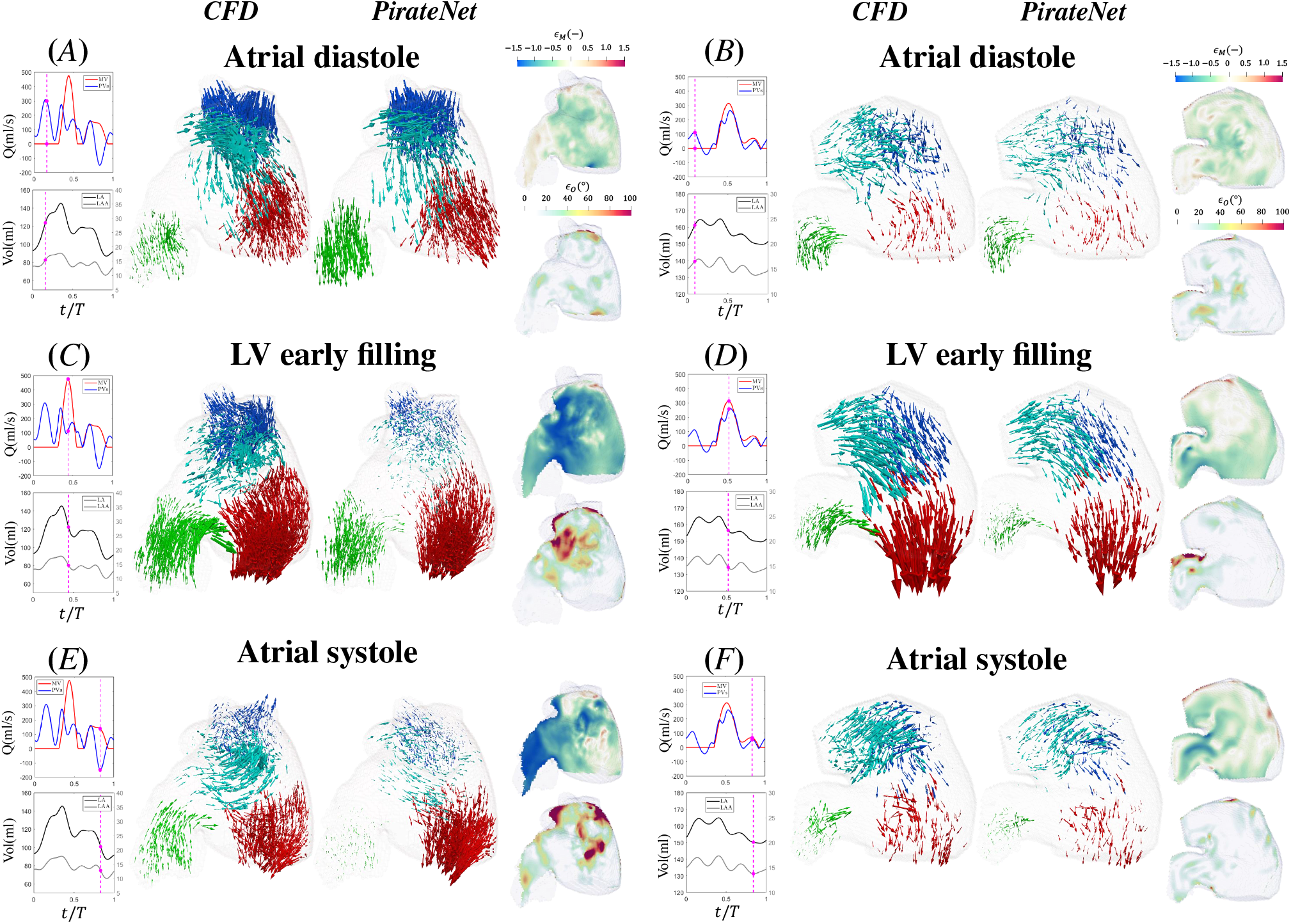
Instantaneous 3D velocity vector fields computed by CFD and inferred by contrast-trained PirateNet, together with normalized errors in PirateNet-inferred velocity magnitude (*ϵ*_*M*_) and angular orientation (*ϵ*_*O*_). Left panels (*A,C,E*) show case 3 (LAAT/TIA-negative, normal LA function), whereas right panels (*B,D,F*) show case 5 (LAAT/TIA-positive, impaired LA function). Each row represents one phases of the cardiac cycle: Top panels (*A, B*) represent atrial diastole; center panels (*C, D*) represent early LV filling; bottom panels (*E, F*) represent and atrial systole. As temporal reference, flow-rate waveforms through the pulmonary veins and mitral valve, together with LA and LAA volume traces, are shown alongside the vector fields; magenta dashed lines indicate the corresponding time instants. Vectors are colored by proximity to anatomical regions: right PVs (blue), left PVs (cyan), LAA (green), and MV (red). To facilitate visualization, the length of PINN-inferred velocity vectors are scaled by a factor of 2.0 in case 3 and 1.8 in case 5, respectively.

During atrial diastole (Fig. 6*A*–*B*), the PINN-inferred velocity field agreed well with the CFD reference in both patients, reproducing the PV inflow jets, their collision in the LA body, and the subsequent redirection of flow toward the MV and the LAA. The associated LAA filling pattern was also captured. The *ϵ*_*M*_ map indicated a mild underestimation of velocity magnitude in the LA body (*ϵ*_*M*_ *<* 0), with a slight overestimation in parts of the LAA (*ϵ*_*M*_ *>* 0). Additionally, the *ϵ*_*O*_ map showed small angular differences between CFD and PINN velocity vectors across most of the LA, indicating good agreement in flow orientation.

Blood continues to enter through the PVs during early LV filling, and LV suction together with passive LA recoil cause the MV to open and drive a strong transmitral jet. During this phase, the LAA also empties, generating outward flow that is partially redirected toward the MV near the LAA ostium. These dominant flow patterns were reproduced by the PINN (Fig. 6*C*–*D*). However, the *ϵ*_*M*_ map revealed a bias toward underestimating velocity magnitudes, particularly in case 3. Nevertheless, flow orientations were largely preserved: elevated *ϵ*_*O*_ values were mainly confined to regions of near-zero velocity (e.g., near the left PVs in case 3 and near the superior LAA wall in case 5).

During atrial systole, the LA contracts, pumping blood into the LV while also generating backflow into the PVs. Accordingly, the LAA continues to evacuate blood through the ostium, forming a jet that is partially deflected toward the MV (Fig. 6*E*–*F*). These dominant flow patterns were reproduced by the PINN, although the LAA emptying jet strength was underestimated in case 3 (consistent with the *ϵ*_*M*_ *<* 0 region in Fig. 6*E*). As in atrial diastole and early LV filling, flow orientation was overall well captured, with elevated *ϵ*_*O*_ values mainly confined to low-speed regions where the velocity direction is effectively irrelevant.

Instantaneous 3D velocity vector fields for the remaining two LAAT/TIA-negative cases are shown in Fig. SI2, while the other two LAAT/TIA-positive cases are shown in Fig. SI3. Overall, these data confirm the trends reported for cases 3 and 5, suggesting that the PINN-inferred flow fields captured the orientation of the CFD ground-truth velocity, but tended to underestimate its magnitude. The globally averaged *ϵ*_*M*_ and *ϵ*_*O*_ for all the cases, shown in Table 2, indicate that the PINN reproduced the flow velocity field particularly well in the LAA, where 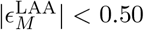 and 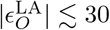 deg for three out of the six cases studied, while *ϵ*_*M*_ increased in the LA body with mixed performances in terms of *ϵ*_*O*_.

## 4 Discussion

This study investigates methods for inferring 4D (3D in space, time-resolved) residence time (*T*_*R*_) fields in a flowing fluid from 4D distributions of a passively advected scalar (*c*) within the same flow. The work is motivated by the existence of medical imaging modalities which enable the quantification of passive scalar dynamics, e.g., 4D CT contrast agent transport, but are incapable of directly measuring residence time distributions. We evaluate on two methods: (i) a physics-informed Neural Network (PINN) constrained by the Navier–Stokes and transport equations for *c* and *T*_*R*_, and (ii) an indicator dilution theory (IDT) formulation that estimates *T*_*R*_ voxel-wise from contrast dynamics curves. Both methods are benchmarked against patient-specific CFD simulations in anatomically realistic, moving left atrial geometries.

Residence time is a relevant biomarker that reflects blood stasis, a central driver of thrombogenesis. We focus on the left atrium (LA), and particularly on the left atrial appendage (LAA), because the LAA is a major site of thrombus formation and embolic stroke. In non-valvular atrial fibrillation (AF), approximately 90% of left atrial thrombi originate in the LAA [5], and AF itself accounts for ~20–30% of all ischemic strokes. The LAA also contributes to thromboembolic risk in the absence of documented AF: 15–38% of atrial thrombi in non-AF cardiomyopathy arise from the LAA.

Patient-specific CFD studies can quantify LAA residence time and related hemodynamic stasis metrics [19–23]. However, routine clinical translation of CFD remains limited by the need for detailed patient-specific boundary conditions, meticulous geometric meshing, and substantial computational resources [30]. Color-Doppler ultrasound remains the first-line modality for cardiac flow imaging, but transesophageal echocardiography—the technique used to visualize the LAA—is invasive, time-consuming, and provides only limited assessment of 3D flow. 4D flow MRI enables direct measurement of 3D flow patterns and stasis [6–8], but its spatiotemporal resolution in the LAA is generally insufficient. CT angiography provides high spatial resolution and has shown that impaired LAA opacification or delayed contrast washout is associated with stroke risk and LAA thrombus formation [12, 13]. Taken together, these constraints underscore the need for imaging-based approaches that can reliably infer residence time in the LAA.

Recent CT studies have linked impaired LAA opacification and delayed washout to LAA thrombus formation and stroke risk [13, 14]. Building on this premise, Severance *et al*. showed that IDT can be applied to clinically acquired multi-cycle 4D contrast CT to generate 3D maps of LAA transport parameters, including an asymptotic residence time *T*_*R*_(**x**, *t* →∞), using voxelwise gamma-variate modeling [37]. In IDT, a downstream concentration–time curve is modeled as the convolution of an input bolus with a transport (impulse-response) function [60]. Historically, this framework has been used in reduced-order, compartment-based models to estimate global indices such as flow, mean transit time, and effective volume [38, 62]. In modern contrast-enhanced CT and MRI, closely related IDT formulations are applied at the voxel level in tissue perfusion imaging, where each voxel is treated as an effective vascularized compartment and deconvolution or parametric modeling yields voxelwise perfusion metrics [63–65].

Here, we extend IDT to a moving intracardiac chamber by constructing a PV-referenced impulse response for each chamber voxel and combining it with a tank-in-series model along flow pathlines to obtain voxelwise, time-resolved residence-time maps *T*_*R*_(**x**, *t*) from contrast dynamics. This PV-referenced formulation isolates the intrinsic chamber response from upstream transport history avoiding subtracting large, nearly equal upstream transit-time estimates (e.g., ostium- and LAA-based fits), which can be poorly conditioned when the chamber contribution is small compared with total venous-to-atrial transit. Additionally, by computing *T*_*R*_(**x**, *t*) rather than the limit *T*_*R*_(**x**, *t* →∞), our estimate is naturally tied to the observed number of cardiac cycles and reduces the risk of overestimation from extrapolating an unobserved washout tail. We expect this pathline-based method should translate to other convection-dominated vascular domains, e.g., intracranial aneurysms, carotid disease, or abdominal aortic aneurysms.

PINNs have emerged as a powerful framework to merge measurement data with physical laws in fluid mechanics and related areas of computational science [36], with applications ranging from aerodynamics to cardiovascular flows [66]. In cardiovascular biomechanics, PINNs have been used to infer latent hemodynamic variables—including pressure fields, wall shear stress, and constitutive parameters—from sparse or indirect measurements [67–70]. A canonical benchmark setting in which PINNs can infer latent velocity fields is advection of a passive scalar, because scalar dynamics provide dense spatiotemporal supervision while velocity remains unobserved. This has motivated tracer-/contrast-informed PINN formulations for flow and parameter inference in biological and medical-imaging contexts [71–73].

A substantial fraction of cardiovascular PINN studies has emphasized reduced-order hemodynamic models and/or vascular domains with relatively simple geometries and little or no wall motion. By comparison, applications targeting fully 3D intracardiac flow in atria or ventricles remain relatively scarce despite recent advances [57, 74]. Consistent with this gap, recovering intracardiac flow fields from contrast-enhanced CT remains underexplored in the PINN literature. Explicit inference of blood residence time is even less commonly considered in cardiovascular PINN studies.

Several aspects of our PINN formulation are tailored to the specific challenges of intracardiac flow reconstruction from contrast data. First, the use of spatiotemporal Fourier feature embeddings helps alleviate the spectral bias of standard MLPs and improves the representation of high-frequency flow features [51]. This is particularly important for capturing sharp inflow jets and complex LAA recirculation patterns. Second, we enforce the initial condition of residence time as a hard constraint by reparameterizing the *T*_*R*_ output [52], ensuring that *T*_*R*_ is exactly zero at (*t* = *t*_0_) without requiring an additional loss term. Third, we frame the learning problem in non-dimensional variables scaled by clinically meaningful quantities such as MV diameter and peak E-wave velocity, which improves numerical conditioning and may facilitate comparison across patients with different hemodynamic regimes. Lastly, we use PirateNet structure through adaptive residual connections, reported to help stabilize the training and mitigate initialization difficulties in training deep PINNs [47]. PirateNet yielded improved accuracy in *T*_*R*_ inference, especially when the vanilla MLP-based structure performed poorly. Together, these design choices enable the PINN to reconstruct volumetric fields in a moving LA, where the wall motion is implicitly encoded through the spatiotemporal distribution of contrast.

When compared with CFD, both PINN- and IDT-inferred *T*_*R*_ fields reproduce the expected trends: higher residence time in the LAA than in the LA body, especially in distal LAA regions, and systematically higher *T*_*R*_ in LAAT/TIA-positive cases than in negative ones. Beyond these qualitative trends, IDT shows the closest agreement with CFD across the full range of *T*_*R*_ values, capturing sharp spatial gradients (e.g., at the LAA ostium) with only a mild tendency to overestimate the upper tail. The PINN also recovers the main *T*_*R*_ patterns, but produces smoother maps and tends to underestimate high-*T*_*R*_ regions, most noticeably in distal LAA. Despite these biases, both methods preserve patient ranking and separate LAAT/TIA-positive from LAAT/TIA-negative cases based on mean LAA *T*_*R*_.

The opposite biases of IDT and PINN reflect the different approximations each method makes. IDT models transport with no explicit diffusion, which can overestimate *T*_*R*_. By contrast, the PINN balances data fit with PDE residuals, favoring smooth, low-frequency solutions in advection-dominated settings due to spectral bias [36, 75]. This phenomenon can smear sharp features and compress the high-*T*_*R*_ tail, leading to underestimation of the largest residence times.

In addition to residence time, the PINN reproduces the main atrial flow structures across the cardiac cycle, including PV inflow jets, their collision and redirection toward the MV, as well as LAA filling and recirculation. PINN also captures the phase-dependent flow reorganization associated with MV opening and closing: during early LV filling it recovers the dominant transmitral jet and the concurrent LAA emptying flow through the ostium, and during atrial systole it reproduces the LAA evacuation jet and the associated flow redirection toward the MV. Across cases, the agreement is stronger in flow orientation than in magnitude, with the PINN underestimating peak speeds while preserving the overall vector field topology. This behavior is consistent with the global velocity statistics.

A key contribution of this study is showing that contrast dynamics can support two complementary inference pathways: a PINN that reconstructs full-field hemodynamics (including residence time) and an IDT-based approach that yields residence-time maps in seconds from contrast curves alone. The PINN provides velocity and pressure fields together with *T*_*R*_, whereas IDT trades that detail for speed and robustness and is therefore appealing for clinical translation. This complementarity suggests a tiered workflow: IDT for rapid stasis screening, and the PINN for cases where detailed flow information is needed. Beyond this, rapid IDT inference could guide a second-stage (nested) PINN. For example, the IDT-inferred residence time *T*_*R*,IDT_(**x**, *t*) could be treated as an additional “measurement” by adding a data-mismatch term, e.g., ℒ_IDT_ = |*T*_*R*,PINN_ − *T*_*R*,IDT_|^2^, and/or it could be included as an extra network input so the PINN learns fields conditioned on an IDT-based stasis estimate (with physics enforced using material derivatives parameterized by *T*_*R*,IDT_). These hybrid IDT–PINN strategies could improve PINN accuracy in high-*T*_*R*_ regions and reduce underestimation of peak velocities, while preserving the PINN’s ability to recover velocity orientation.

Iodinated CT contrast can, in principle, change the local density and viscosity of blood [76], creating both a practical modeling issue and a more fundamental concern: the injection could perturb the very hemodynamics we aim to infer from the contrast dynamics. In our setting, however, pilot CFD simulations at clinically relevant iodine concentrations (Boussinesq approximation) did not show noticeable changes in left atrial velocity fields, suggesting that this effect is negligible for the protocols considered here. If such effects were non-negligible, they could be incorporated into the PINN by modifying the momentum/transport physics terms to allow property variations (e.g., variable density/viscosity), whereas the IDT approach does not require an explicit flow model and would not need to be modified beyond treating the measured curves as the effective transport response.

Training the PINN remains computationally demanding (on the order of 10–15 hours per patient on a high-end GPU), and retraining is currently required for each new patient, although this burden could be reduced with transfer learning (i.e., warm-start PINNs; [77]). Furthermore, our validation relies on CFD simulations, which themselves depend on assumptions regarding blood rheology, wall motion, and boundary conditions [20, 27–29, 78], and are not equivalent to *in vivo* measurements. A natural next step is therefore to leverage emerging multi-cycle 4D contrast CT datasets [37] to build matched CFD and PINN models directly from clinically acquired contrast data, and to compare CFD-derived *T*_*R*_ against *T*_*R*_ inferred from the CT time curves using IDT and PINNs. Ultimately, direct validation remains challenging because no clinical modality can measure intracardiac residence time as a ground-truth field.

The IDT formulation, while computationally efficient, assumes that the LA behaves as a linear, time-invariant system with respect to contrast input. In our derivation, this approximation is justified by analyzing transport at a fixed cardiac phase and treating the phase-locked velocity field as “steady on the mean” across cycles, with negligible molecular diffusion (high Péclet number). Under these conditions, the phase-stacked voxel time curves can be interpreted through an effective impulse response (modeled here with a tank-in-series/gamma-variate form). In reality, LA contraction, PV inflow variability, and beat-to-beat fluctuations may violate these assumptions, particularly in AF. Moreover, IDT does not provide velocity or pressure fields and thus cannot capture mechanistic features such as swirling patterns, shear rate, or pressure gradients.

Despite our encouraging results, there are clear paths to improve both accuracy and practicality. On the modeling side, performance may benefit from more expressive architectures (e.g., operator-learning or multi-network decompositions), adaptive or curriculum-based loss weighting, and hard constraints for inlet residence-time conditions [52]. On the implementation side, migrating to modern platforms (e.g., TensorFlow 2.x or JAX with XLA compilation) could substantially reduce training time and ease deployment [79]. On the data side, an important next step is to train and test on clinically acquired 4D contrast CT with fewer cardiac phases, motion artifacts [80], and variable injection protocols.

## 5 Conclusion

In summary, we present and validate two complementary, physics-based approaches to infer left atrial stasis from contrast dynamics: a PINN that reconstructs full flow and residence time maps, and an IDT framework that produces residence time maps rapidly from contrast curves alone. Benchmarked against patient-specific CFD in moving atrial geometries, both methods recover key LAA stasis patterns and differentiate LAAT/TIA-positive from LAAT/TIA-negative cases based on mean LAA *T*_*R*_. Together, these results support contrast-enhanced 4D CT as a scalable route toward quantitative, patient-specific assessment of LAA thrombosis risk.

## Acknowledgments

This study was supported by the US National Institutes of Health (NIH) under award numbers R01HL160024 (to JCA, EMV, AK) and R01HL158667 (to JCA), the Spanish Research Agency under award number PID2023-146861OB-I00 (to PML) and the Instituto de Salud Carlos III, Spain, under grant number PI21/00274-PACER1 (to JB). The EU—European Regional Development Fund also supported this work.

## Supporting Information

**Figure SI1:**
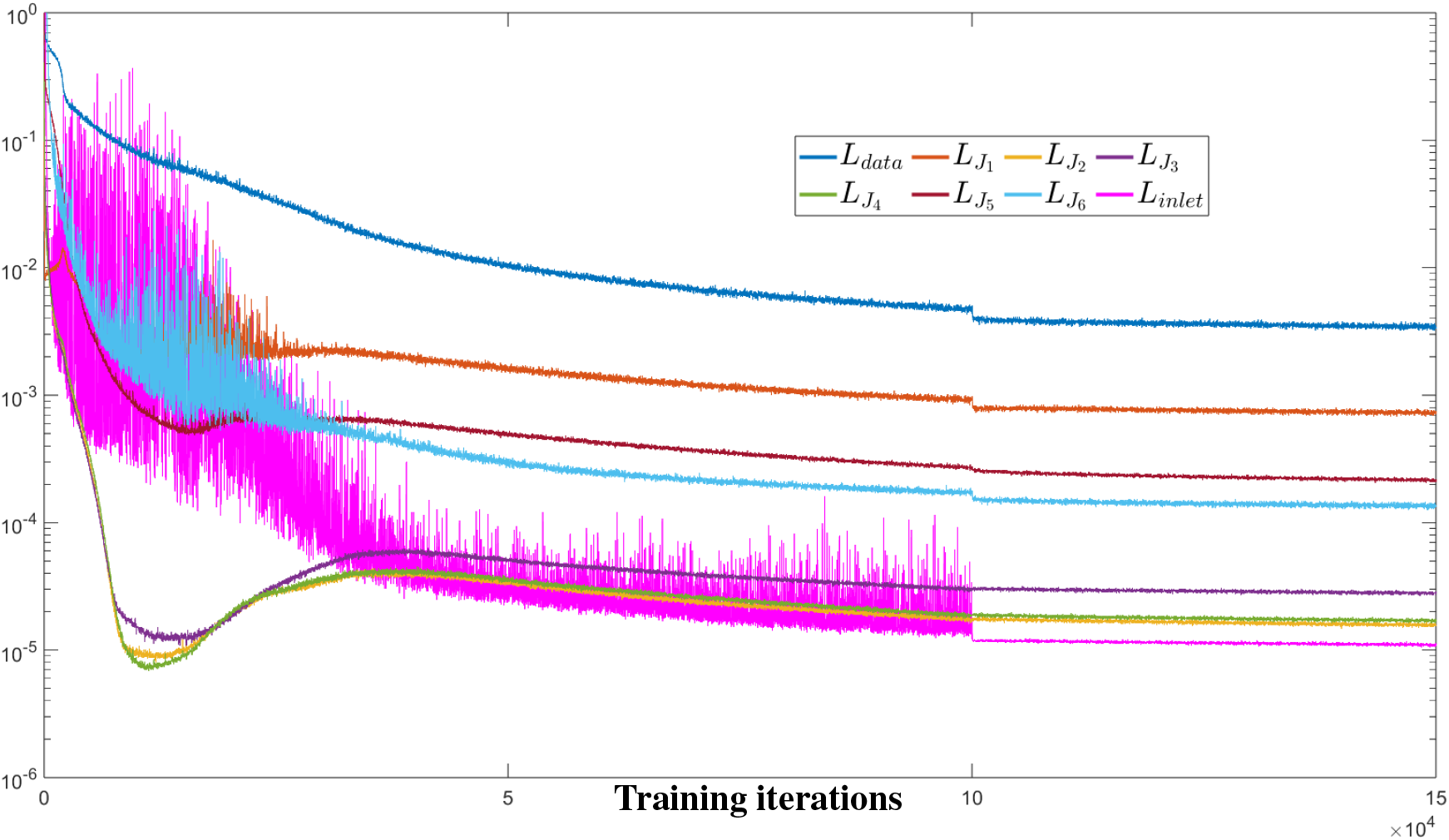
Loss curves for each loss component of a representative case 1 from PirateNet training.

**Figure SI2:**
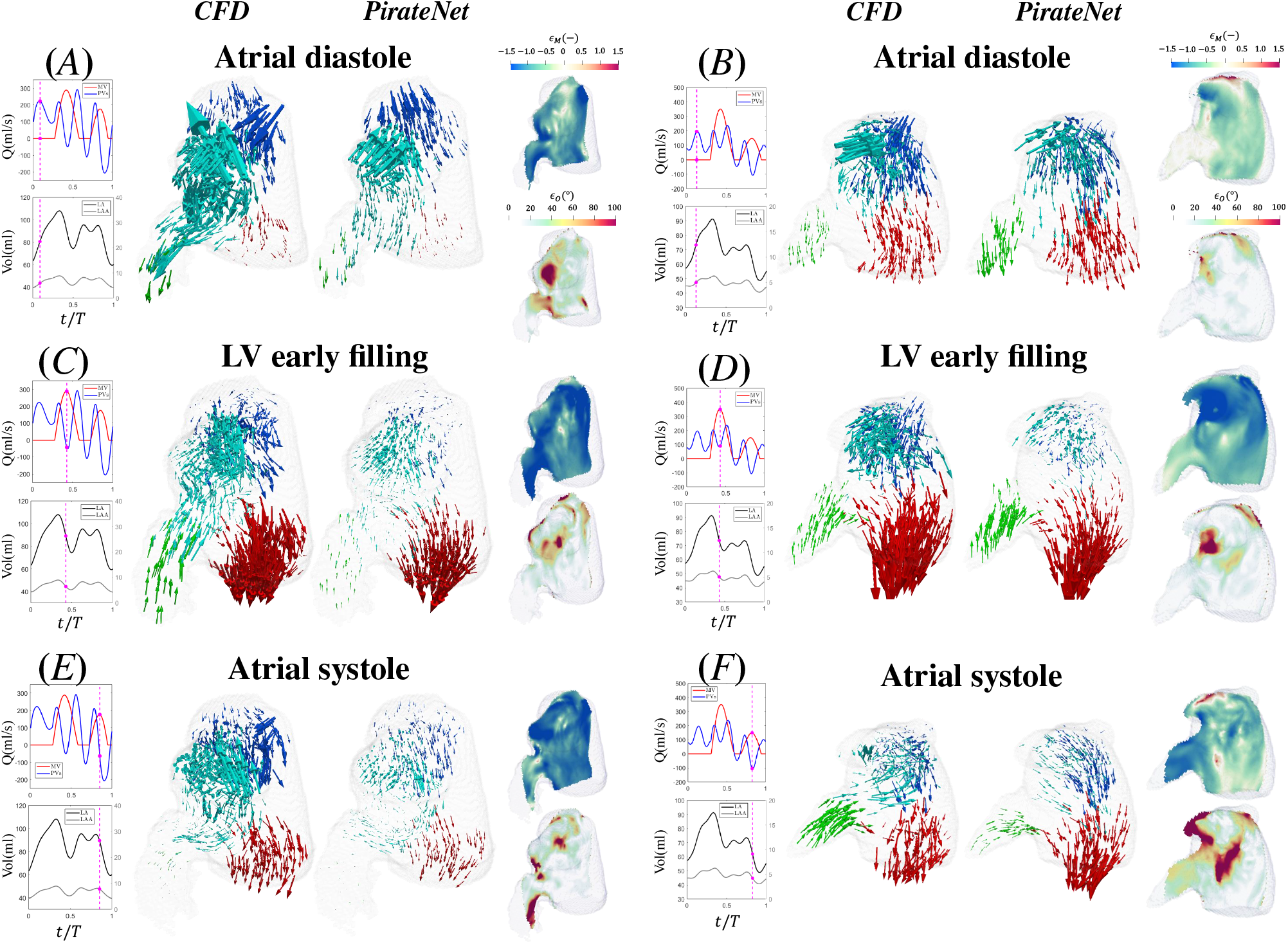
Instantaneous 3D velocity vector fields computed by CFD and inferred by contrast-trained PirateNet, together with normalized errors in PirateNet-inferred velocity magnitude (*ϵ*_*M*_) and angular orientation (*ϵ*_*O*_). Left panels (*A,C,E*) show case 1 (LAAT/TIA-negative, normal LA function), whereas right panels (*B,D,F*) show case 2 (LAAT/TIA-negative, normal LA function). Each row represents one phases of the cardiac cycle: Top panels (*A, B*) represent atrial diastole; center panels (*C, D*) represent early LV filling; bottom panels (*E, F*) represent and atrial systole. As temporal reference, flow-rate waveforms through the pulmonary veins and mitral valve, together with LA and LAA volume traces, are shown alongside the vector fields; magenta dashed lines indicate the corresponding time instants. Vectors are colored by proximity to anatomical regions: right PVs (blue), left PVs (cyan), LAA (green), and MV (red). To facilitate visualization, the length of PINN-inferred velocity vectors are scaled by a factor of 2.5 in case 1 and 2.0 in case 2, respectively.

**Figure SI3:**
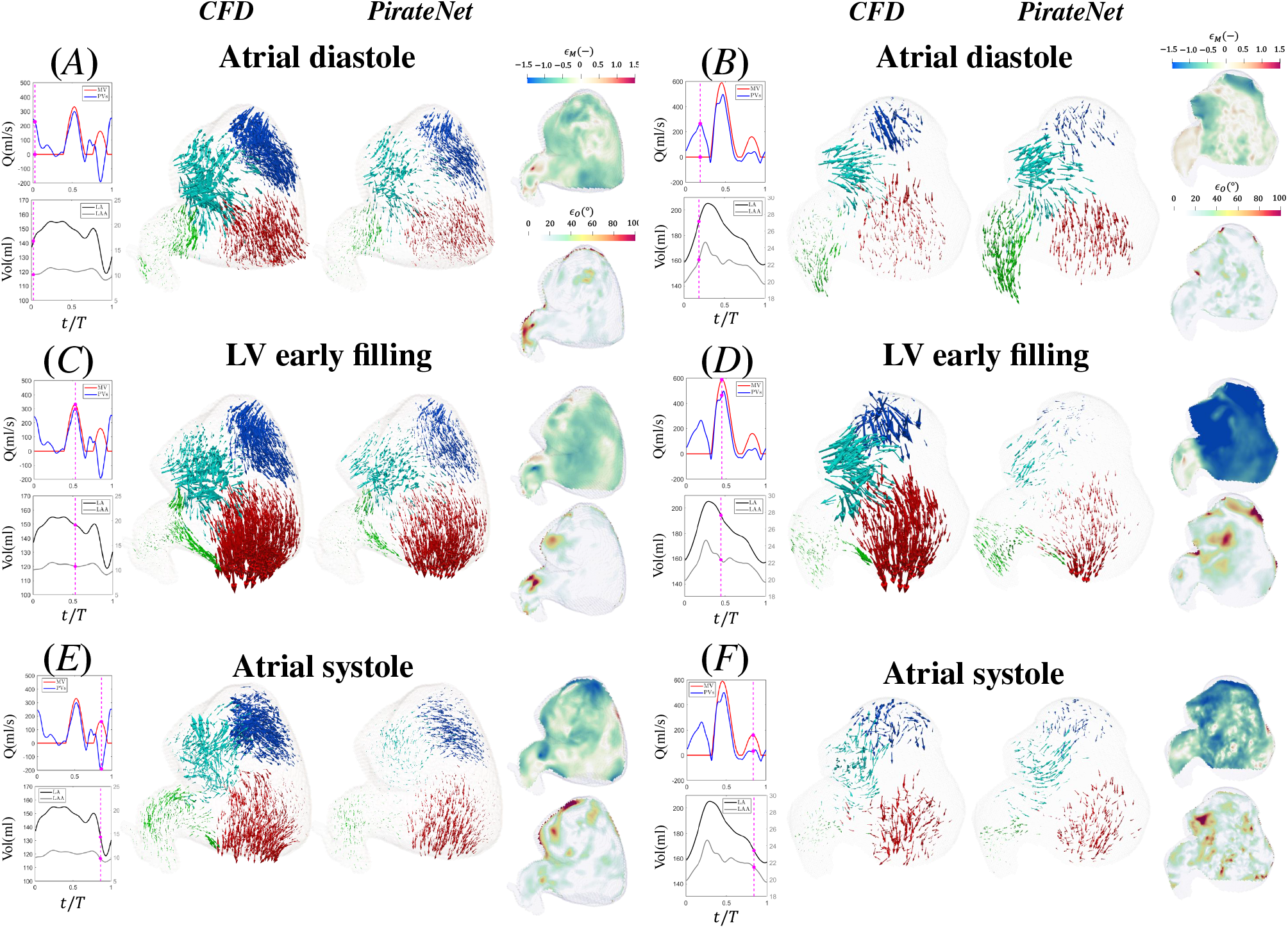
Instantaneous 3D velocity vector fields computed by CFD and inferred by contrast-trained PirateNet, together with normalized errors in PirateNet-inferred velocity magnitude (*ϵ*_*M*_) and angular orientation (*ϵ*_*O*_). Left panels (*A,C,E*) show case 4 (LAAT/TIA-positive, impaired LA function), whereas right panels (*B,D,F*) show case 6 (LAAT/TIA-positive, impaired LA function). Each row represents one phases of the cardiac cycle: Top panels (*A, B*) represent atrial diastole; center panels (*C, D*) represent early LV filling; bottom panels (*E, F*) represent and atrial systole. As temporal reference, flow-rate waveforms through the pulmonary veins and mitral valve, together with LA and LAA volume traces, are shown alongside the vector fields; magenta dashed lines indicate the corresponding time instants. Vectors are colored by proximity to anatomical regions: right PVs (blue), left PVs (cyan), LAA (green), and MV (red). To facilitate visualization, the length of PINN-inferred velocity vectors are scaled by a factor of 2.0 for both cases.

**Figure SI4:**
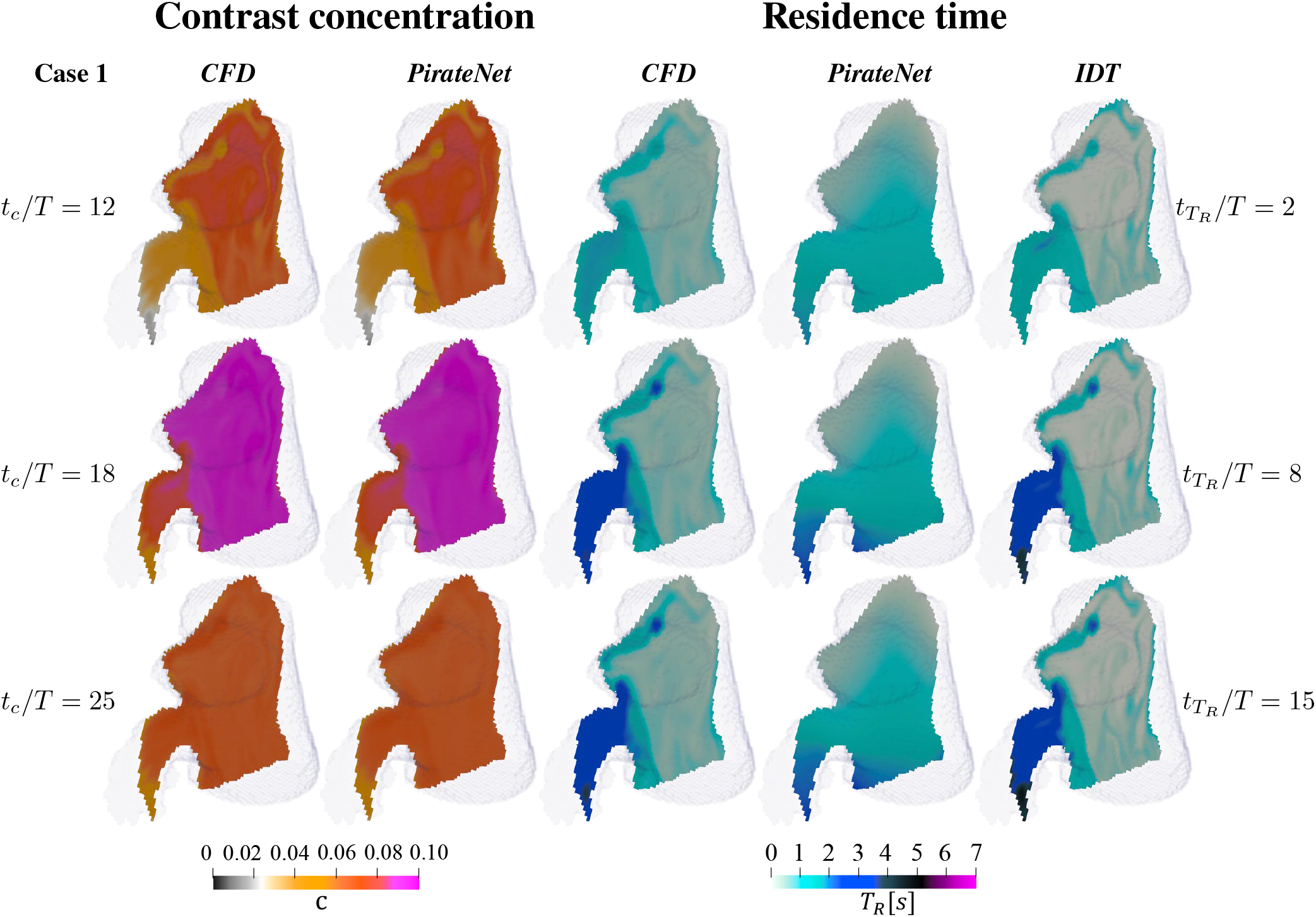
Contrast concentration (*c*) and residence time (*T*_*R*_) maps comparison of case 1 between CFD, PirateNet, and IDT. Three 2D sections of the left atrium are shown at selected time instants. Contrast concentration snapshots are shown at *t*_*c*_*/T* = 12, 18, and 25, whereas residence time snapshots are shown at 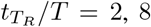, and 15. Here *t*_*c*_ is referenced to the contrast-training window (cycles 10–25), and 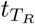 is referenced to the start of residence time integration. Contrast concentration is dimensionless, whereas residence time is given in seconds.

**Figure SI5:**
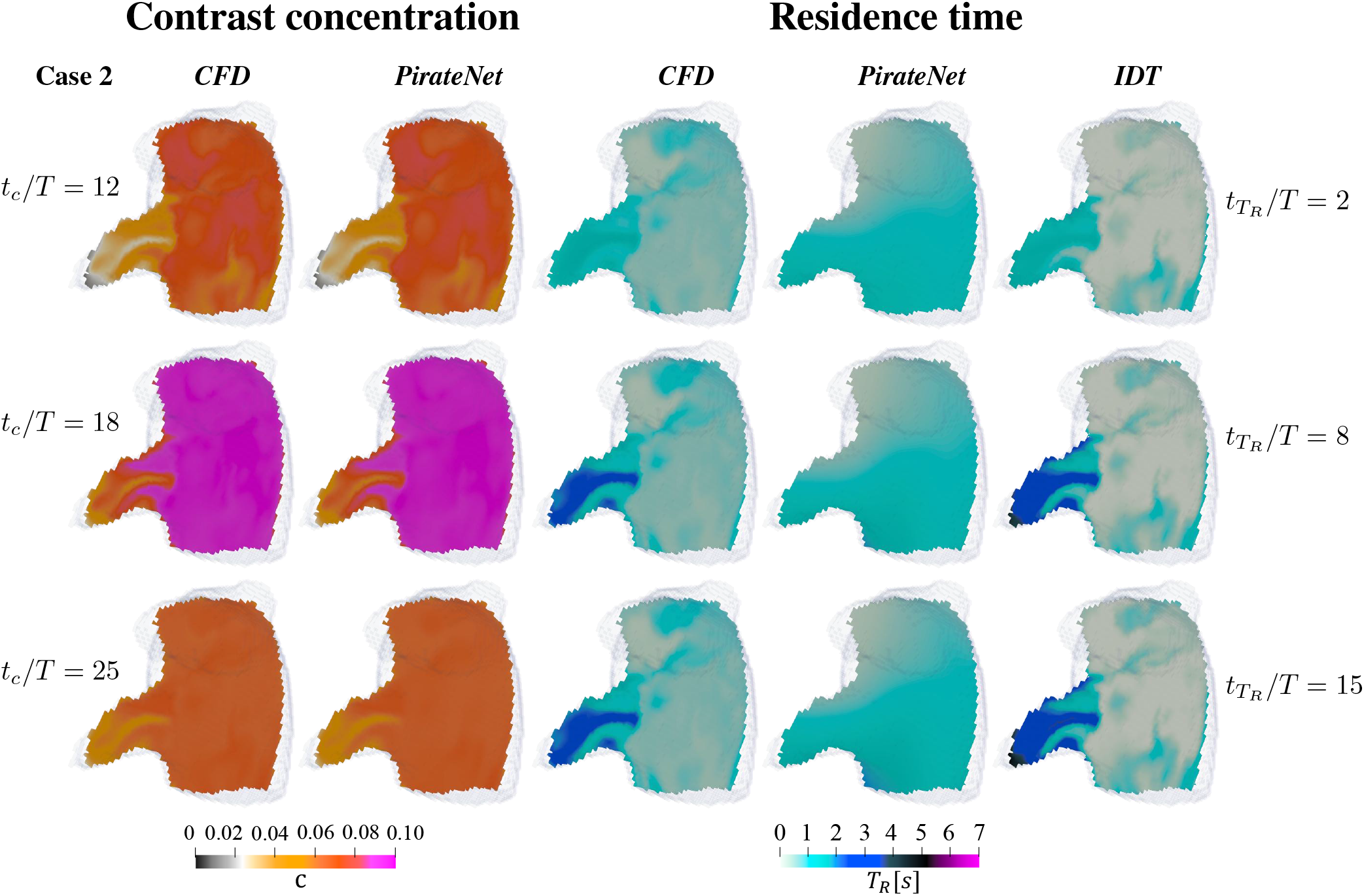
Contrast concentration (*c*) and residence time (*T*_*R*_) maps comparison of case 2 between CFD, PirateNet, and IDT. Three 2D sections of the left atrium are shown at selected time instants. Contrast concentration snapshots are shown at *t*_*c*_*/T* = 12, 18, and 25, whereas residence time snapshots are shown at 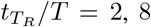, and 15. Here *t*_*c*_ is referenced to the contrast-training window (cycles 10–25), and 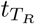 is referenced to the start of residence time integration. Contrast concentration is dimensionless, whereas residence time is given in seconds.

**Figure SI6:**
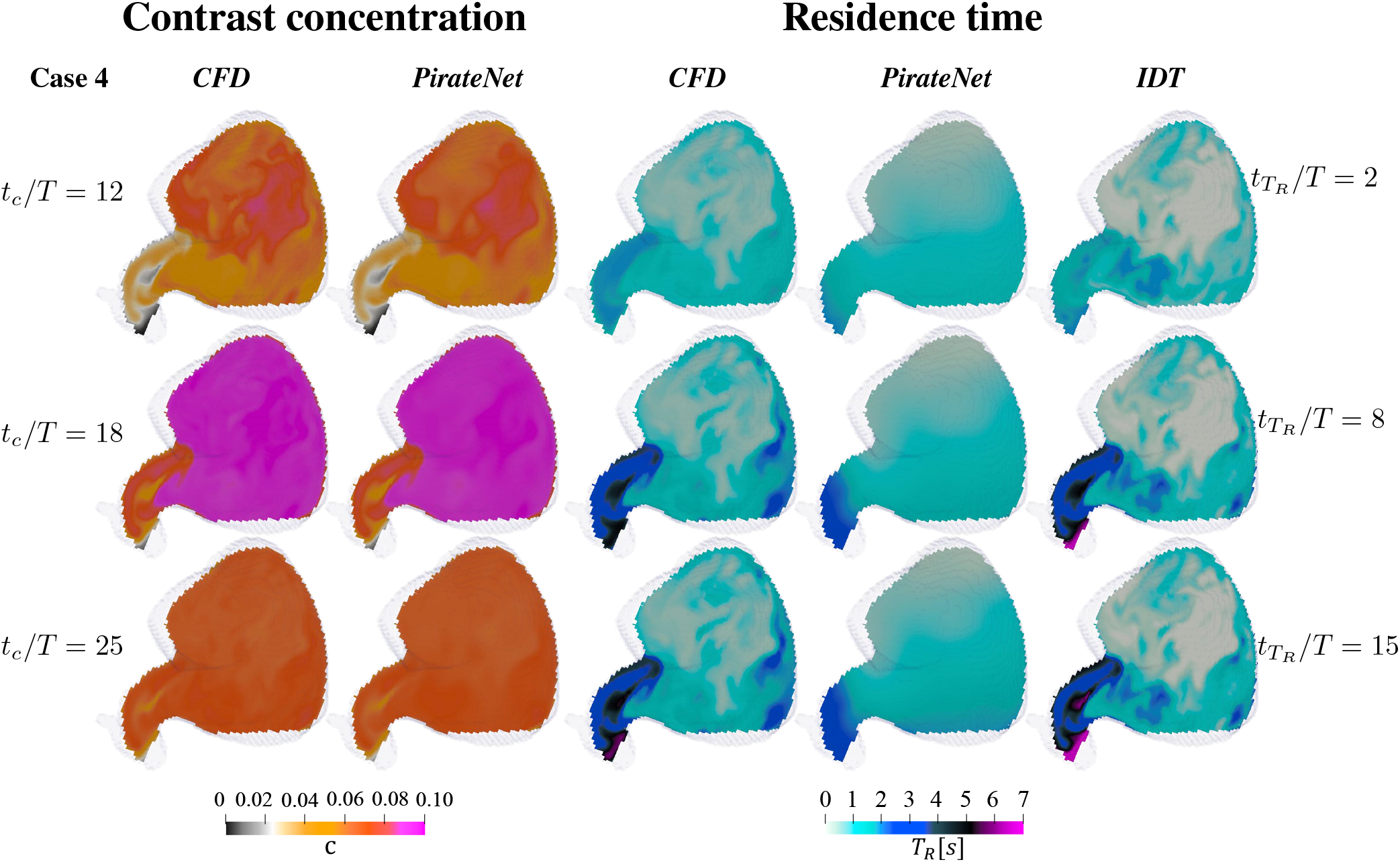
Contrast concentration (*c*) and residence time (*T*_*R*_) maps comparison of case 4 between CFD, PirateNet, and IDT. Three 2D sections of the left atrium are shown at selected time instants. Contrast concentration snapshots are shown at *t*_*c*_*/T* = 12, 18, and 25, whereas residence time snapshots are shown at 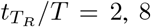, and 15. Here *t*_*c*_ is referenced to the contrast-training window (cycles 10–25), and 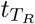 is referenced to the start of residence time integration. Contrast concentration is dimensionless, whereas residence time is given in seconds.

**Figure SI7:**
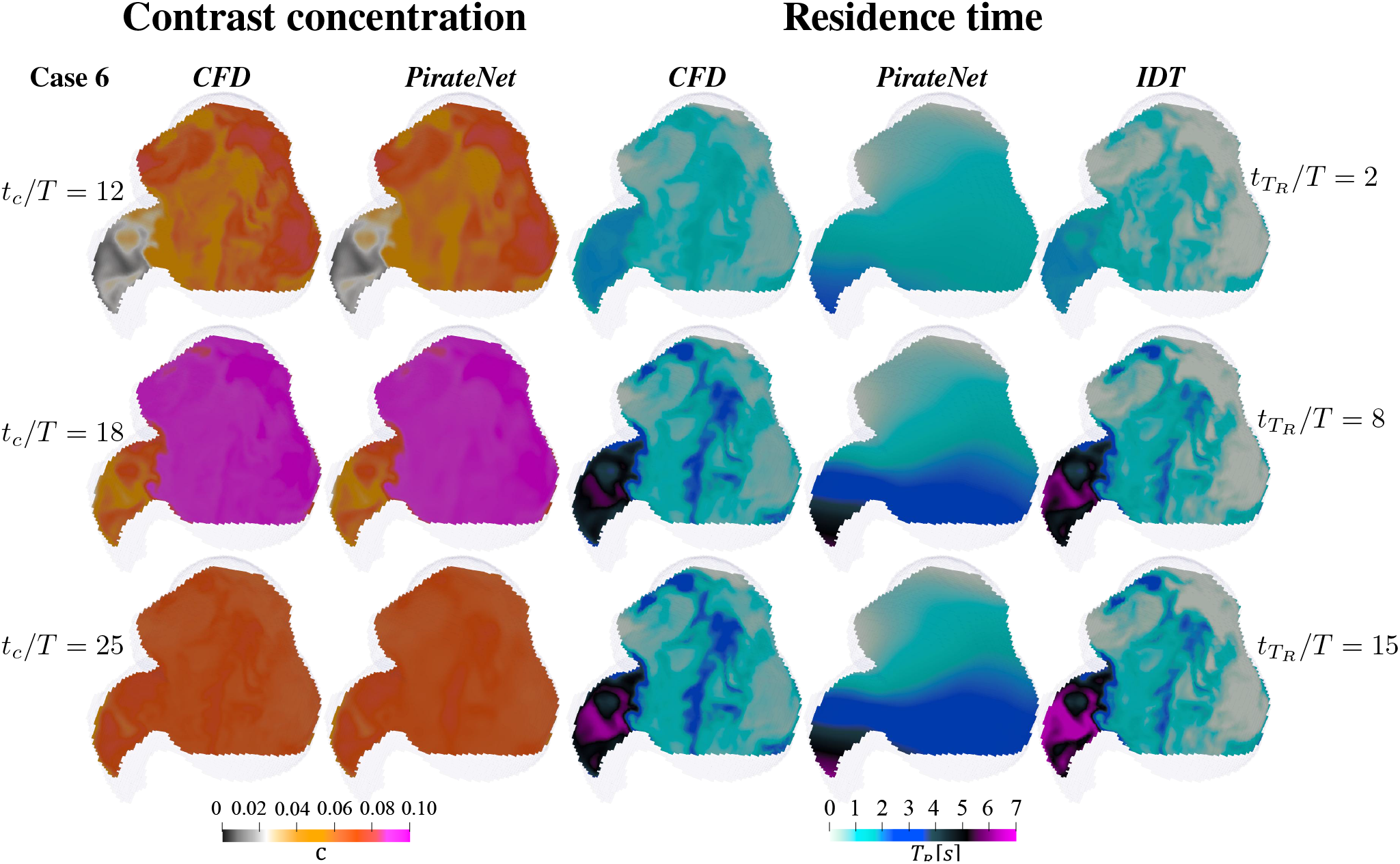
Contrast concentration (*c*) and residence time (*T*_*R*_) maps comparison of case 6 between CFD, PirateNet, and IDT. Three 2D sections of the left atrium are shown at selected time instants. Contrast concentration snapshots are shown at *t*_*c*_*/T* = 12, 18, and 25, whereas residence time snapshots are shown at 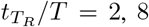, and 15. Here *t*_*c*_ is referenced to the contrast-training window (cycles 10–25), and 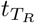 is referenced to the start of residence time integration. Contrast concentration is dimensionless, whereas residence time is given in seconds.

**Figure SI8:**
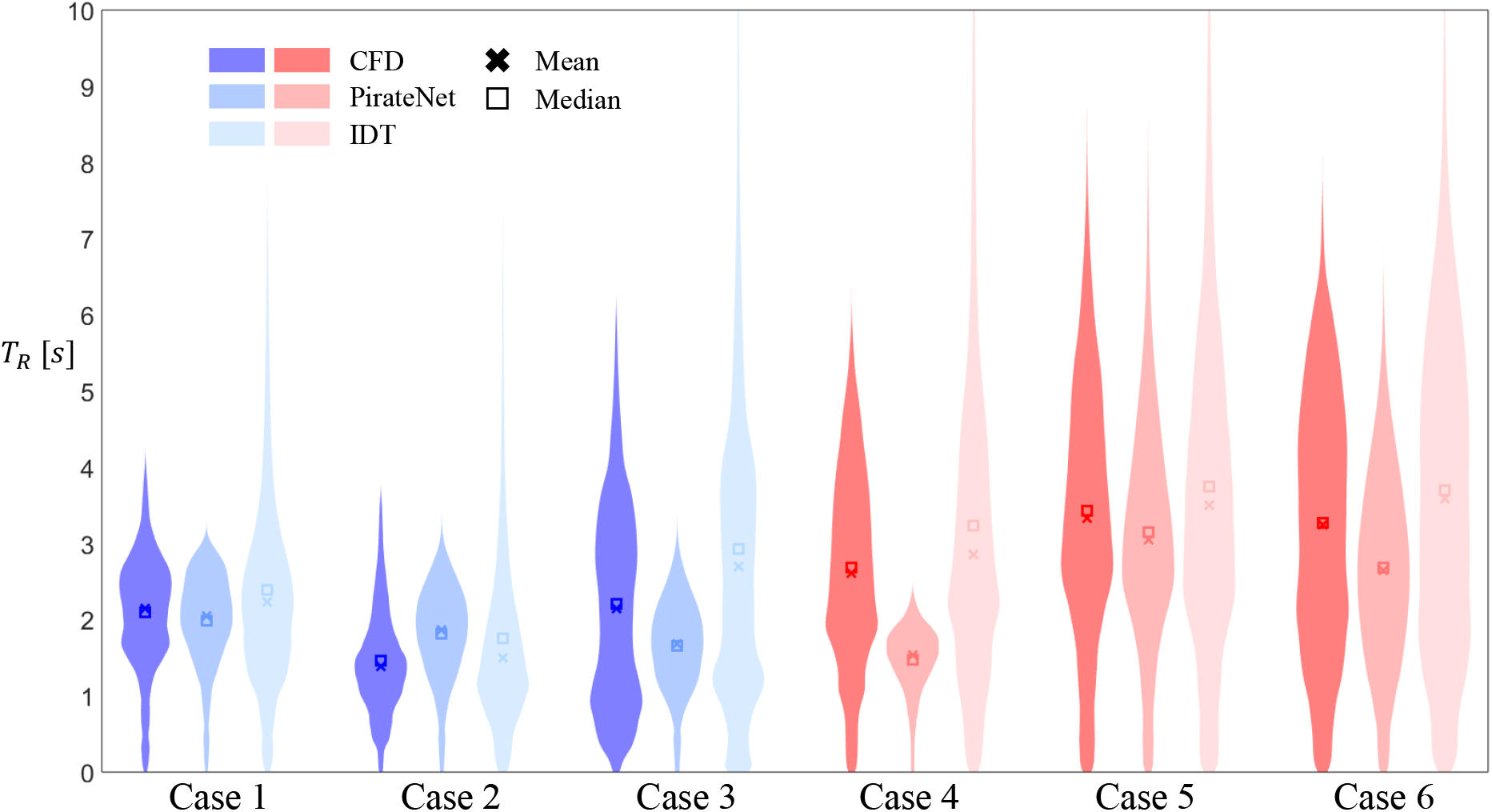
Violin plots of the probability density function with mean and median of the residence time *T*_*R*_ from CFD, PirateNet, and IDT inside the left atrial appendage are shown for the *N* = 6 studied cases.

